# β-Adrenergic inhibition of exocytotic surface deposition of MHCII molecules in reactive astrocytes is mediated by amisyn

**DOI:** 10.64898/2026.08.19.745707

**Authors:** Julijan Vršnik, Mićo Božić, Zara Bunc, Maja Potokar, Keita Sugiyama, Klemen Dolinar, Sergej Pirkmajer, Gregor Anderluh, Marko Kreft, Ira Milošević, Jernej Jorgačevski, Robert Zorec, Matjaž Stenovec

**Affiliations:** Laboratory of Neuroendocrinology-Molecular Cell Physiology, Institute of Pathophysiology, Faculty of Medicine, University of Ljubljana, Zaloška 4, 1000 Ljubljana, Slovenia; Celica Biomedical, Tehnološki park 24, 1000 Ljubljana, Slovenia; Faculty of Chemistry and Chemical Technology, Večna pot 113, 1000 Ljubljana, Slovenia; Department of Pharmacology, Kurume University School of Medicine, 67 Asahi-machi, Kurume 830-0011, Japan; Laboratory for Molecular Neurobiology, Institute of Pathophysiology, Faculty of Medicine, University of Ljubljana, Zaloška 4, 1000 Ljubljana, Slovenia; Department of Molecular Biology and Nanobiotechnology, National Institute of Chemistry, Hajdrihova 19, 1000 Ljubljana, Slovenia; Biotechnical Faculty, University of Ljubljana, Jamnikarjeva 101, Ljubljana, Slovenia; NIHR Oxford Biomedical Research Centre, Nuffield Department of Medicine, Wellcome Centre for Human Genetics, University of Oxford, Roosevelt Drive, Oxford, OX3 7BN, UK

## Abstract

Degeneration of the locus coeruleus, a noradrenergic nucleus, reduces noradrenaline bioavailability in the central nervous system and promotes neuroinflammation via reactive astrocytes, although the underlying mechanisms remain unclear. We investigated whether interferon-γ-induced expression of major histocompatibility complex class II (MHCII), a marker of pro-inflammatory reactive astrocytes, is regulated by adrenergic receptors and amisyn. β-Adrenergic, but not α-adrenergic, stimulation increased cyclic adenosine monophosphate (cAMP) and reduced MHCII expression, as detected immunocytochemically, in human and rat astrocytes. β-Adrenergic treatment altered transient exocytosis of lysosome-like vesicles, increasing event frequency and reducing fusion-pore conductance and dwell time, thereby limiting MHCII surface expression. Overexpression of wild-type amisyn inhibited surface expression of MHCII and the lysosomal marker CD63 and reduced fusion-pore conductance and dwell time. Conversely, amisyn knockdown enhanced full fusion exocytosis of larger vesicles and abolished β-adrenergic effects, indicating that amisyn mediates β-adrenergic inhibition of exocytosis and MHCII surface deposition.

**Graphical abstract:** 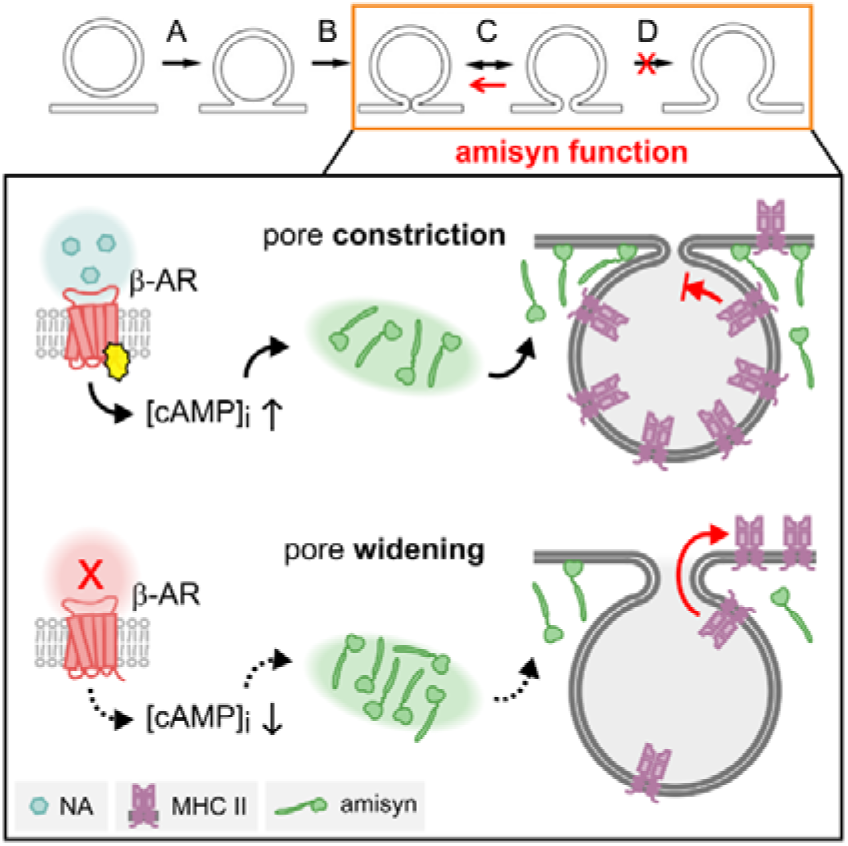

**Top:** States in which a vesicle interacts with the plasma membrane during exocytosis: A, hemifusion; B, formation of a narrow fusion pore; C, transient fusion with reversible pore opening; D, full fusion with irreversible pore expansion. Amisyn promotes a narrow fusion pore (C) and hinders the progression to state D. **Bottom:** Mechanism of fusion pore regulation by β-adrenergic receptor (β-AR) and cyclic AMP; amisyn redistributes to the inner face of the plasmalemma, keeping the fusion pore constricted and reducing the probability of the fusion pore entering the full-fusion stage, thus inhibiting the density of surface-expressed MHCII antigen-presenting molecules. NA, noradrenaline.

**Main points:**

- Loss of noradrenergic tone after locus coeruleus degeneration promotes astrocyte-mediated neuroinflammation.
- β-Adrenergic, but not α-adrenergic, stimulation increases cAMP and reduces MHCII expression in interferon γ-induced reactive astrocytes.
- β-Adrenergic signalling limits MHCII surface expression by reducing fusion-pore conductance and dwell time during transient vesicle fusion.
- Amisyn mediates this β-adrenergic inhibition; its knockdown enhances full vesicle fusion and abolishes β-adrenergic effects.

## Introduction

In many neurodegenerative diseases, including Alzheimer disease and Parkinson disease, the locus coeruleus, the principal noradrenaline-releasing nucleus in the central nervous system (CNS), undergoes degeneration^1–3^. This reduces the bioavailability of noradrenaline (NA)^4^, impairing a key modulator of synaptic plasticity that acts through astrocytes^5^ and contributes to broader astrocyte dysfunction, including altered regulation of neuroinflammation^6^.

Astrocytes maintain homeostasis in the CNS and support tissue defence^7^ through reactive astrogliosis, a process involving morphological, molecular, and functional remodelling in response to injury, disease, or infection^8^. When dysregulated, reactive astrogliosis can suppress homeostatic astrocyte functions and promote or accelerate neuroinflammation,^6,9^ contributing to neurodegenerative diseases^10–13^ and autoimmune disorders^14^. Understanding the cellular mechanisms that limit reactive astrogliosis, particularly how NA inhibits astrocyte-mediated neuroinflammation, is therefore essential^15^.

Impaired noradrenergic signalling has been linked to immunosuppression, primarily through increased intracellular concentration of cyclic AMP ([cAMP]_i_), which reduces antigen presentation and cytokine secretion in macrophages, dendritic cells, and facultative or tissue-resident antigen-presenting cells, including astrocytes and Langerhans cells^16^. This pathway also regulates gene expression, including CIITA, the major histocompatibility complex class II (MHCII) transactivator^17^ and MHCII surface expression, likely through mechanisms involving vesicle trafficking and vesicle–plasma membrane interactions in astrocytes^18^ and dendritic cells^19^.

One possible mechanism underlying noradrenergic inhibition of antigen presentation is a reduced ability of vesicles carrying antigens and antigen presentation machinery to fuse with the plasma membrane and/or release their cargo upon fusion. These mechanisms can be examined at the single-vesicle level using membrane capacitance recordings, which directly report changes in plasma membrane area during vesicle fusion^20,21^, while signalling pathways and proteins that regulate exocytotic vesicle–plasma membrane merger are manipulated.

A promising candidate regulator is amisyn (syntaxin-binding protein 6 [STXBP6]), a brain-enriched protein that negatively regulates exocytosis and is expressed in astrocytes (ProteinAtlas, accessed on 10 May 2023; networkglia, accessed on 22 December 2021). Amisyn has been implicated in diabetes^22^ and neurodevelopmental disorders, including autism spectrum disorder^23^, and developmental epileptic encephalopathy^24^. It coimmunoprecipitates with syntaxin-1a and syntaxin-4^25^, is recruited to vesicle fusion sites in a cAMP-dependent manner^26^, reduces exocytotic discharge from chromaffin and PC12 cells (a pheochromocytoma-derived cell line of the rat adrenal medulla)^27^ and may regulate fusion-pore stability, as shown by amperometry studies^27,28^.

We asked whether amisyn participates in NA-mediated regulation of lysosome-associated antigen presentation^18^, a process induced by pro-inflammatory cytokines that promote reactive astrogliosis^29^. Astrocytes were exposed to the cytokine, interferon γ (IFNγ), which activates the IFNγ-R1 receptor and the Janus kinase/signal transducer and activator of transcription protein pathway, triggering transcriptional responses^30^ across multiple genes^31,32^, and robust upregulation of MHCII expression in astrocytes^32,33^, a hallmark of reactive astrocytes^18^. In summary, β-adrenergic treatment attenuates MHCII surface expression by suppressing large-vesicle exocytosis, reducing fusion-pore conductance and dwell time, and promoting amisyn recruitment to the plasma membrane. Amisyn overexpression mimics these effects, whereas amisyn knockdown enhances vesicle exocytosis and abolishes β-adrenergic inhibition. Thus, loss of NA in neurodegeneration may promote astrocyte-mediated neuroinflammation by reducing amisyn-dependent β-adrenergic suppression of antigen presentation.

## Results

### IFN**γ**-induced MHCII expression in human and rat astrocytes is inhibited by noradrenaline

IFNγ-induced expression of antigen-presenting MHCII molecules was observed in human astrocytes (Fig. 1a, top left), consistent with recent findings in human reactive astrocytes^34^. Similarly, IFNγ treatment increased MHCII expression in primary rat astrocytes (Fig. 1a, bottom left,), confirming previous observations^18,35^. The addition of noradrenaline (NA) resulted in a significant reduction in MHCII expression in both astrocyte populations (Fig. 1a, top and bottom right; Fig. 1b, c; Extended Data Fig. 1). These results indicate that NA inhibits antigen presentation in rat and human astrocytes, consistent with early flow cytometric studies in IFNγ-treated astrocytes, which showed dose-dependent inhibition of MHCII surface expression by NA^36^.

**Fig. 1.**
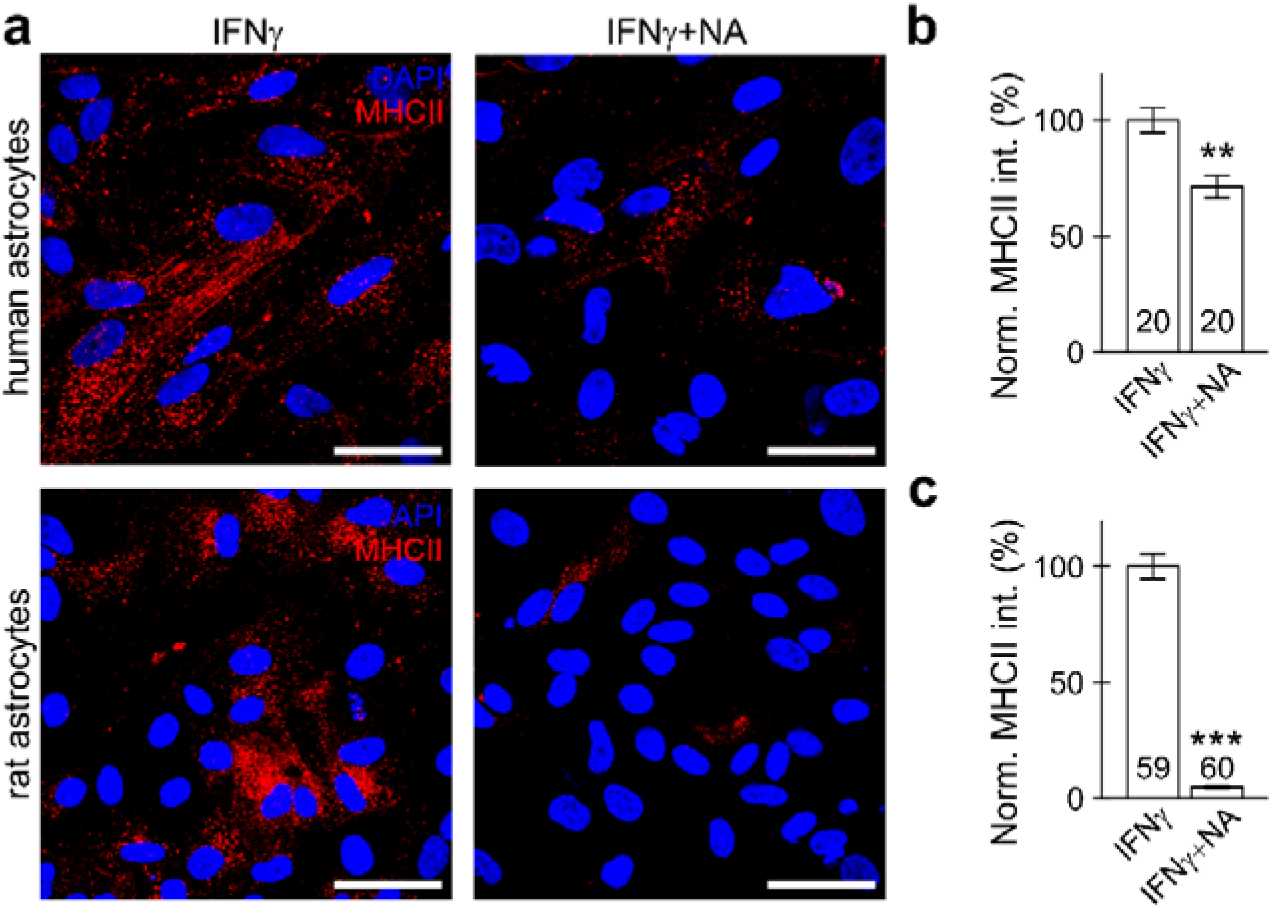
Noradrenaline (NA) reduces cellular MHCII expression in interferon γ (IFNγ)-treated human and rat astrocytes. (**a**) Confocal images of fixed and permeabilized human (top) and rat (bottom) astrocytes immunolabelled with the anti-MHCII antibody and the corresponding fluorescent secondary antibody (MHCII, red), and by the fluorescent nuclear dye DAPI (blue), after 48 h treatment with 100 U/ml of IFNγ (left), and co-treatment with NA (100 µM; right). (**b, c**) Graphs displaying mean (±SEM) normalized (norm.) MHCII fluorescence per cell (relative to IFNγ treatment) subjected to (co)treatment with IFNγ and NA in a primary cell culture of human (**b**) and rat (**c**) astrocytes. Numbers at the bottom of the bars indicate the number of images analysed; \*\**P* < 0.01, \*\*\**P* < 0.001 (ANOVA on ranks followed by Dunn’s test).

### Surface expression of MHCII in astrocytes is inhibited by **β**-adrenergic signalling

We next examined rat astrocytes, in which NA produced the strongest inhibitory effect (Fig. 1). In reactive astrocytes, MHCII localizes to lysosome-like vesicles and is delivered to the plasma membrane^18^. Surface MHCII was therefore analysed in live, non-permeabilized cells using an anti-MHCII antibody.

IFNγ treatment (60 U/ml, 48 h) induced surface MHCII expression in most, but not all, astrocytes (Fig. 2a). Co-treatment with NA (100 µM) significantly reduced the proportion of MHCII-positive (MHCII+) cells (Fig. 2b). Combining phenylephrine (PE), an α-adrenergic agonist, with the β-adrenergic antagonist propranolol (Pro), did not reduce MHCII labelling (Fig. 2c). In contrast, the β-adrenergic agonist isoprenaline (Iso), combined with the α-adrenergic antagonist terazosin (Trz), nearly abolished surface MHCII expression (Fig. 2d). The membrane-permeable cAMP analogue, dibutyryl cyclic adenosine monophosphate (dbcAMP; 1 mM), produced a similar inhibitory effect, implicating β-adrenergic receptor–cAMP signalling in NA-mediated suppression of MHCII surface expression (Fig. 2e). Comparable results were obtained in fixed, permeabilized cells (Extended Data Fig. 1).

**Fig. 2.**
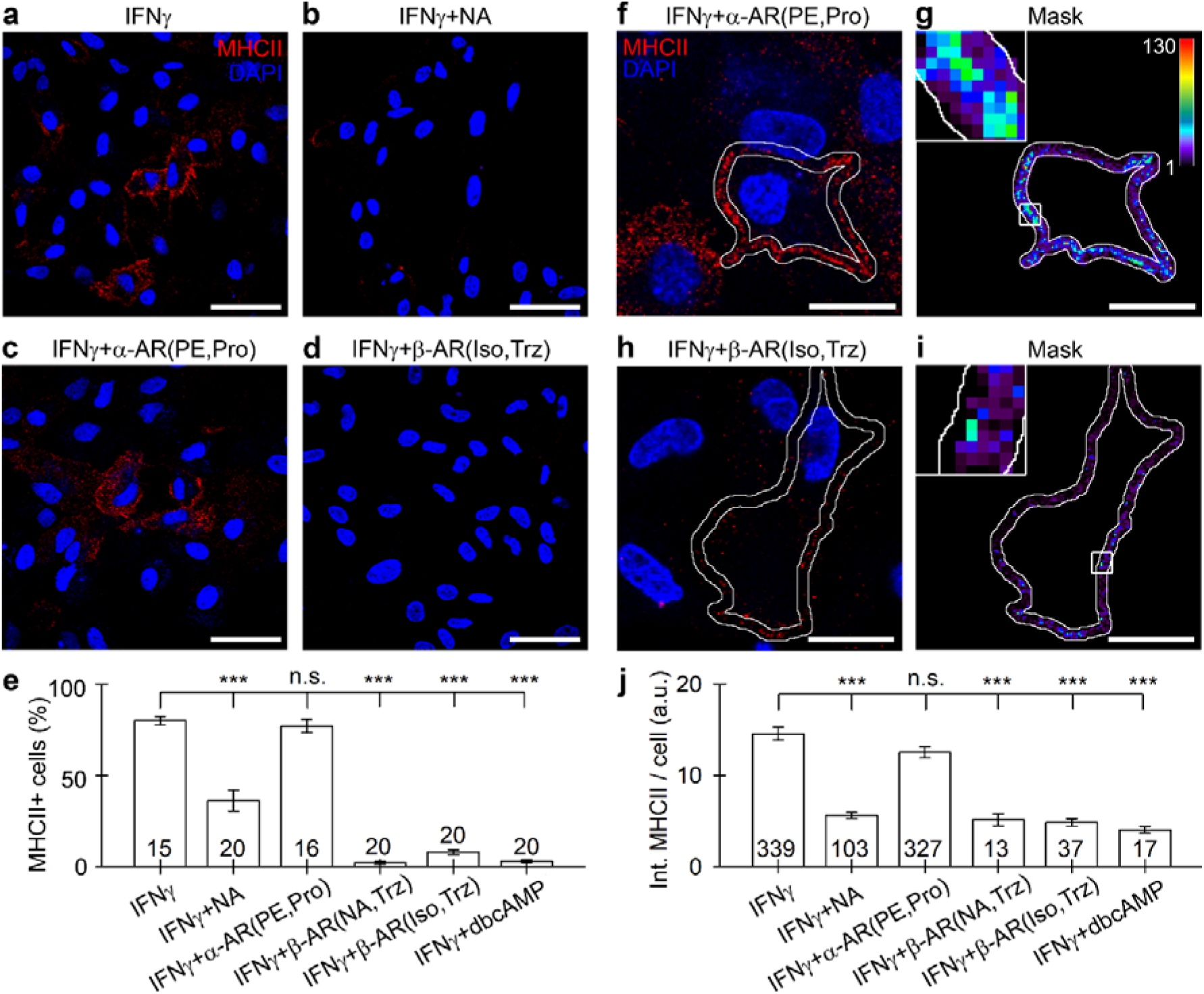
Treatment with β-adrenergic agonists or dibutyryl cyclic adenosine monophosphate (dbcAMP) reduces the percentage of MHCII+ astrocytes and MHCII surface expression in interferon γ (IFNγ)-treated astrocytes. (**a–d**) Confocal images of live, non-permeabilized astrocytes immunolabelled by the anti-MHCII antibody and corresponding fluorescent secondary antibody (MHCII, red), and by the fluorescent nuclear dye (DAPI; blue), acquired in culture treated with 60 U/ml of IFNγ for 48 h (**a**); co-treated with 100 µM noradrenaline (NA) (**b**); co-treated with IFNγ, the α-adrenergic agonist 100 µM phenylephrine (PE), and 2 µM β-adrenergic antagonist propranolol (Pro) (**c**); co-treated with IFNγ, the β-adrenergic agonist 100 µM isoprenaline (Iso), and 2 µM α-adrenergic antagonist terazosin (Trz) (**d**). Scale bars: 50 µm. (**e**) Graph displaying percentage of MHCII+ astrocytes (MHCII) after different co-treatments with IFNγ, α- and β-adrenergic (ant)agonists or dbcAMP. Note that activation of β-, but not α-adrenergic receptors (ARs), substantially reduced the percentage of MHCII+ astrocytes. Numbers at the bottom of the bars indicate the number of images analysed (statistical unit). The total numbers of cells analysed for each treatment were 427, 259, 431, 645, 666, 710, respectively (from left to right). This experiment was independently repeated on three different cell cultures. \*\*\**P* < 0.001; n.s. not significant (ANOVA on ranks followed by Dunn’s test). (**f, h**) Confocal images of non-permeabilized astrocytes immunolabelled by the anti-MHCII antibody and the corresponding fluorescent secondary antibody (MHCII, red), and by the fluorescent nuclear stain DAPI (blue) after 48 h co-treatment with 60 U/ml of IFNγ, 100 µM PE and 2 µM Pro (**f**); or co-treatment with 60 U/ml of IFNγ, 100 µM Iso and 2 µM Trz (**h**). Note the relatively high MHCII immunofluorescence in astrocytes co-treated with IFNγ, PE and Pro (f) in comparison with astrocytes co-treated with IFNγ, Iso and Trz (H). (**g, i**) Pseudocoloured display of MHCII immunofluorescence obtained from (**f**) and (**h**); MHCII fluorescence intensity is displayed as an intensity scale from 0 to 130 a.u. in pseudocolour (**g**, top right). The insets in (**g**) and (**i**) are magnified (6×) views of MHCII immunofluorescence in a selected region of the astrocyte surface (white rectangle). Scale bars (**f–i**): 20 μm (large images). (**j**) Graph displaying mean (±SEM) MHCII surface expression (fluorescence intensity per cell) in astrocytes subjected to different (co)treatments with IFNγ and α- and β-adrenergic (ant)agonists or dbcAMP. Activation of β-, but not α-ARs, reduced the surface expression (density) of MHCII in astrocytes. Numbers at the bottom of the bars indicate the number of astrocytes analysed (statistical unit). \*\*\**P* < 0.001, n.s. not significant (ANOVA on ranks followed by Dunn’s test).

Peripheral MHCII fluorescence was quantified within a 2-µm band along the cell margin (Extended Data Fig. 2; Fig. 2f–i^18^). IFNγ increased the proportion of MHCII+ astrocytes to 80% ± 2% (*n* = 427), an effect not inhibited via specific α-adrenergic stimulation (IFNγ plus PE and Pro; 77% ± 4%; *n* = 431). NA reduced this proportion to 36% ± 6% (*n* = 259, *P* < 0.001), whereas IFNγ combined with NA plus Trz, Iso plus Trz, or dbcAMP reduced it to 2% ± 1% (*n* = 645), 8% ± 1% (*n* = 467), and 3% ± 1% (*n* = 710), respectively (all *P* < 0.001; ANOVA on ranks with Dunn’s test; Fig. 2e). These findings show that β-, but not α-adrenergic signalling inhibits MHCII surface expression in reactive astrocytes.

### β-Adrenergic stimulation suppresses transient fusion of larger vesicles and reduces fusion-pore size and dwell time in IFNγ-treated astrocytes

Because β-adrenergic stimulation reduced MHCII surface expression (Fig. 2), we examined whether cAMP-dependent regulation involves vesicle delivery, exocytotic insertion, or endocytotic retrieval of MHCII^35,37^. High-resolution membrane capacitance (*C*_m_) measurements were used to monitor single-vesicle fusion and fusion-pore properties^20,21^.

We analysed 211 astrocytes from more than 30 independent primary cultures for a total recording time of 27.5 h (Table 1a). Two types of exocytotic events were detected: transient and full fusion; transient events accounted for approximately 80% of all events (Fig. 3a; Extended Data Fig. 3), consistent with previous reports^20,38,39^. Transient events consisted of at least one discrete increase in *C*_m_ (on step) followed by a step decrease in *C*_m_ (off step) of similar amplitude within 15 s, representing transient single-vesicle fusion with the plasmalemma, reflecting fusion-pore opening and closure, whereas full-fusion events appeared as persistent increases in *C*_m_. Transient (reversible) fusion predominated under all treatment conditions.

**Fig. 3.**
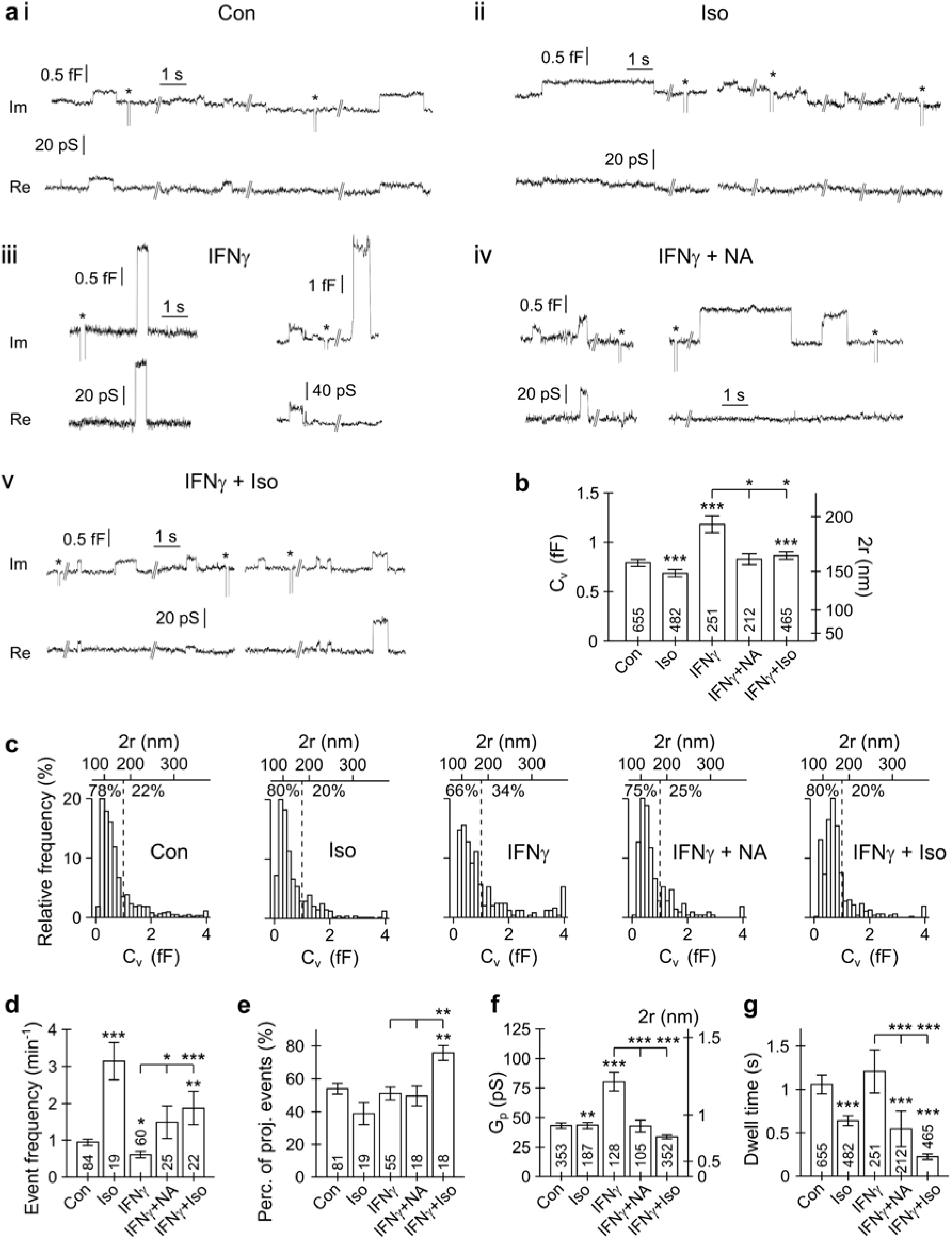
β-Adrenergic stimulation regulates exocytotic fusion-pore properties and reduces IFNγ-induced transient exocytosis of large diameter vesicles. (**a**) Representative recordings of transient vesicle exocytosis in non-treated controls (**i**), isoprenaline (a β-adrenergic agonist, Iso, 100 µM)-treated astrocytes (**ii**), IFNγ-treated astrocytes (**iii**), and astrocytes co-treated with IFNγ and noradrenaline (NA, 100 µM) (**iv**) or Iso (**v**). Asterisks (*) denote calibration pulses. Note the occurrence of large transient exocytotic vesicles in IFNγ-treated astrocytes, but not in non-treated controls and astrocytes co-treated with IFNγ and β-adrenergic agonists (NA, Iso). (**b**) Graph depicting (mean ± SEM) vesicle capacitance (*C*_v_) and diameter (2r, right ordinate) in vesicles undergoing transient exocytosis. A considerable increase in *C*_v_ in IFNγ-treated astrocytes was inhibited in astrocytes co-treated with IFNγ and β-adrenergic agonists (NA, Iso). Note that Iso-treatment reduced *C*_v_ independently of IFNγ treatment, indicating regulation of elementary exocytotic events via β-adrenergic signalling. (**c**) Relative frequency distributions of vesicle capacitance (*C*_v_, bottom) and vesicle diameter (top) of transient exocytotic events in all groups examined. The dashed vertical line delimits the percentage of vesicles smaller or larger than 1 fF (corresponding to vesicle diameter ∼178 nm). Note an increase in the proportion of larger transient exocytotic vesicles in IFNγ-treated astrocytes (*C*_v_ ≥ 1 fF) that may correspond to transient fusion of endolysosomes carrying MHCII^18^. (**d–g**) Graphs showing (mean ± SEM) event frequency (**d**); the percentage of Re (real part of admittance)-projected events (vesicles establishing a narrow fusion pore^38,81^) (**e**); fusion-pore conductance (*G*_p_) (**f**); fusion-pore dwell time (**g**) of transient exocytotic vesicles. β-Adrenergic stimulation (Iso, IFNγ + Iso, IFNγ + NA) reduced fusion-pore conductance, consistent with fusion-pore constriction and decreased fusion-pore stability indicated by the increased frequency of transient fusion events (interpreted as repeated openings and closings of the fusion pore in vesicles unable to proceed to full fusion) (**d**); decreased open pore dwell time (**g**); decreased fusion-pore size, indicated by an increase in percentage of exocytotic vesicles with a narrow pore (**e**); decreased fusion-pore conductance (**f**). Graphs depicting vesicle capacitance (*C*_v_; **b**) and fusion-pore conductance (*G*_p_; **f**) also display the corresponding vesicle diameter or pore diameter on the ordinate at the right. Numbers at the bottom of the bars denote the number of events (**b, f, g**) or the number of recordings (**d, e**) analysed. The statistical symbols depicted above the bars (**b, d–g**) indicate two sets of comparisons. Asterisks above the individual bars indicate comparisons versus the control group (Con); asterisks above the graphic intervals indicate pairwise comparisons versus the IFNγ group. Pairwise comparisons (Dunn’s tests) were conducted after ANOVA on ranks and were Bonferroni corrected. \**P* < 0.05, \*\**P* < 0.01, \*\*\**P* < 0.001.

**Table 1.** Membrane capacitance analysis of full (Full) and transient (Trans.) exocytotic (Exo) events in non-transfected controls and astrocytes co-treated with interferon γ (IFNγ) and either noradrenaline (NA) or isoprenaline (Iso) (Table 1a) Overview. (Table11**b**) Non-transfected astrocytes: analysis of electrophysiological measurements (mean ± SEM) of transient exocytotic events. Statistical significance versus control (Con): \**P* < 0.05; \*\**P* < 0.01; \*\*\**P* < 0.001 (Dunn’s test). (Table 1c) Pairwise statistical comparisons of transient exocytotic events between experimental groups exposed to different treatments: Con, non- transfected controls; Iso, isoprenaline-treated astrocytes; astrocytes treated with interferon γ (IFNγ); astrocytes co-treated with IFNγ and noradrenaline (IFNγ + NA) or isoprenaline (IFNγ + Iso).

| Table 1a<br>Treatment | Number of cells | Effective time (min) | Total event count | Total event frequency (min<sup>-1</sup>) | Event count |  | Percentage (%) |  | Event frequency (min<sup>-1</sup>) |  |
| --- | --- | --- | --- | --- | --- | --- | --- | --- | --- | --- |
|  |  |  |  |  | Full Exo | Trans. Exo | Full Exo | Trans. Exo | Full Exo | Trans. Exo |
| Con | 84 | 692.0 | 836 | 1.21 | 181 | 655 | 21.7 | 78.3 | 0.26 | 0.95 |
| Iso | 19 | 153.3 | 585 | 3.82 | 103 | 482 | 17.6 | 82.4 | 0.67 | 3.14 |
| IFNγ | 60 | 413.5 | 327 | 0.79 | 76 | 251 | 23.2 | 76.8 | 0.18 | 0.61 |
| IFNγ+NA | 25 | 142.2 | 272 | 1.91 | 60 | 212 | 22.1 | 77.9 | 0.42 | 1.49 |
| IFNγ+Iso | 23 | 248.1 | 602 | 2.43 | 137 | 465 | 22.8 | 77.2 | 0.55 | 1.87 |

IFNγ , interferon $\gamma$ ; Iso, isoprenaline; NA, noradrenaline.
| **Table 1b** | **Number of events** | **Number of cells** | **$C_v$ (fF)** | **Event frequency (events/min)** | **Percentage of projected events (%)** | **$G_p$ (pS)** | **$G_p/C_v$ (pS/fF)** | **Dwell time (s)** |
| --- | --- | --- | --- | --- | --- | --- | --- | --- |
| **Treatment** |  |  |  |  |  |  |  |  |
| Control | 655 | 84 | $0.79 \pm 0.03$ | $0.94 \pm 0.08$ | $53.8 \pm 3.2$ | $43.2 \pm 2.2$ | $44.1 \pm 1.5$ | $1.05 \pm 0.1$ |
| Iso | 482 | 19 | $0.68 \pm 0.03^{***}$ | $3.14 \pm 0.5^{**}$ | $38.7 \pm 6.7$ | $43.3 \pm 2.5^{**}$ | $39.5 \pm 1.6$ | $0.64 \pm 0.1^{***}$ |
| IFNγ | 251 | 60 | $1.17 \pm 0.08^{***}$ | $0.6 \pm 0.08^*$ | $50.9 \pm 3.8$ | $80.3 \pm 7.9^{***}$ | $48.9 \pm 3.1$ | $1.2 \pm 0.24$ |
| IFNγ + NA | 212 | 25 | $0.82 \pm 0.05$ | $1.49 \pm 0.44$ | $49.5 \pm 5.9$ | $42.8 \pm 5$ | $38.3 \pm 2.1$ | $0.54 \pm 0.2^{***}$ |
| IFNγ + Iso | 465 | 22 | $0.86 \pm 0.03^{***}$ | $1.87 \pm 0.45$ | $75.6 \pm 4.5^{**}$ | $33.6 \pm 1.7$ | $39.4 \pm 1.7$ | $0.22 \pm 0.0^{***}$ |

| **Table 1c** | **$C_v$ (fF)** | **Event frequency (events/min)** | **Percentage of projected events (%)** | **$G_p$ (pS)** | **Dwell time (s)** |
| --- | --- | --- | --- | --- | --- |
| **Comparison** |  |  |  |  |  |
| Con - Iso | <0.001*** | <0.001*** | 0.188 | 0.002** | <0.001*** |
| Con - IFNγ | <0.001*** | 0.023* | 1 | <0.001*** | 0.124 |
| Con - IFNγ + NA | 0.172 | 0.226 | 1 | 1 | <0.001*** |
| Con - IFNγ + Iso | <0.001*** | 0.004** | 0.014* | 0.232 | <0.001*** |
| IFNγ - IFNγ + NA | 0.058 | 0.012* | 1 | <0.001*** | 0.018* |
| IFNγ - IFNγ + Iso | >0.5 | <0.001*** | 0.002** | <0.001*** | <0.001*** |

$C_v$ , vesicle capacitance; $G_p$ , pore conductance. Bonferroni-corrected $P$ values result from individual comparisons (Dunn's test, t test) after ANOVA on ranks or one-way ANOVA. \* $P < 0.05$ ; \*\* $P < 0.01$ ; \*\*\* $P < 0.001$ .

In projected transient events, fusion-pore conductance (*G*_p_) and pore diameter were determined from the imaginary (Im) and real (Re) admittance signals (Extended Data Fig. 3biii). Vesicle capacitance (*C*_v_) correlated positively with *G*_p_ in untreated controls (Extended Data Fig. 3c; r = 0.66; *P* < 0.001), as reported previously^40^.

IFNγ increased the mean *C*_v_ of transiently fusing vesicles^18^, whereas Iso reduced it. Mean vesicle diameter increased from 145 ± 3 nm in controls to 173 ± 6 nm after IFNγ treatment (*P* < 0.001), but decreased to 134 ± 4 nm with IFNγ plus NA and to 147 ± 3 nm with IFNγ plus Iso (Fig. 3ai–v, b; Table 1b).

Iso alone reduced the mean diameter to 133 ± 3 nm (Fig. 3b; Table 1b). Frequency distributions showed that these reductions primarily reflected fewer fusion events involving large vesicles (*C*_v_ ≥ 1 fF; diameter ≥ 178 nm; Fig. 3c). Thus, β-adrenergic stimulation preferentially suppresses transient fusion of larger vesicles in IFNγ-reactive astrocytes.

We analysed event frequency, projected events, fusion-pore conductance (*G*_p_), and pore dwell time to determine how β-adrenergic signalling affects transient fusion pores in IFNγ-treated astrocytes (Fig. 3d–g; Table 1b and Table 1c). IFNγ reduced transient-event frequency, whereas β-adrenergic treatment increased it (Fig. 3d). The proportion of projected events, indicative of narrow fusion pores, increased significantly only with IFNγ plus Iso (Fig. 3e).

IFNγ markedly increased *G*_p_ relative to controls (Fig. 3f), likely reflecting fusion of larger vesicles, given the positive correlation between *C*_v_ and *G*_p_ (Extended Data Fig. 3c). Co-treatment with NA or Iso reduced *G*_p_ compared with IFNγ alone, whereas Iso alone had only a modest effect (Fig. 3f). β-Adrenergic treatment also shortened fusion-pore dwell time two- to sixfold compared with controls and IFNγ-treated astrocytes (Fig. 3g). Thus, β-adrenergic signalling constrains the electrical geometry and stability of the fusion pore, although the underlying molecular mechanism remains to be identified.

### Astrocytes express amisyn, a negative regulator of exocytosis

We identified amisyn (STXBP6) as a candidate mediator of β-adrenergic, cAMP-dependent inhibition of MHCII surface expression through fusion-pore constriction. Amisyn is brain-enriched, expressed in astrocytes (mRNA; <u>Protein Atlas</u>), regulates fusion pores, and acts downstream of cAMP signalling^25,27,28,41^.

Immunolabelling of fixed, permeabilized astrocytes showed predominantly diffuse cytosolic amisyn with additional peripheral staining (Extended Data Fig. 4a). Astrocytes overexpressing amisyn-wild-type (wt)-enhanced green fluorescent protein (EGFP) displayed stronger anti-amisyn fluorescence than non-transfected cells, confirming overexpression of the tagged protein in addition to endogenous amisyn.

Amisyn associates with plasmalemmal phosphatidylinositol 4,5-bisphosphonate (PI(4,5)P_2_) through its N-terminal pleckstrin homology (PH) domain^28^. Accordingly, wt amisyn-EGFP accumulated at the cell periphery, whereas a PI(4,5)P_2_-binding-deficient PH mutant did not (Extended Data Fig. 4b–e), confirming PH-domain-dependent plasma membrane localization, informing the assumption of limited functionality of the PH mutant and the decision to use it as an experimental control for further experiments.

### β-Adrenergic signalling recruits wt amisyn to the astrocyte plasmalemma and reduces MHCII and CD63 surface expression

We next examined whether an increase in β-adrenergic [cAMP]_i_ alters the distribution of amisyn. Astrocytes overexpressing amisyn-wt-EGFP were treated for 1 h with extracellular solution (ECS), NA, Iso, or Iso plus Trz. β-Adrenergic stimulation increased peripheral amisyn fluorescence and induced marked morphological changes, quantified by calculating the cell shape factor (CSF; Fig. 4a–e), consistent with a previous report^42^. The plasmalemma-to-cytosol amisyn ratio increased from 0.97 ± 0.01 in controls to 1.06 ± 0.01 with noradrenaline (NA), 1.15 ± 0.02 with Iso, and 1.09 ± 0.01 with Iso plus Trz (*P* < 0.001; Fig. 4e), indicating recruitment of amisyn to the plasma membrane.

**Fig. 4.**
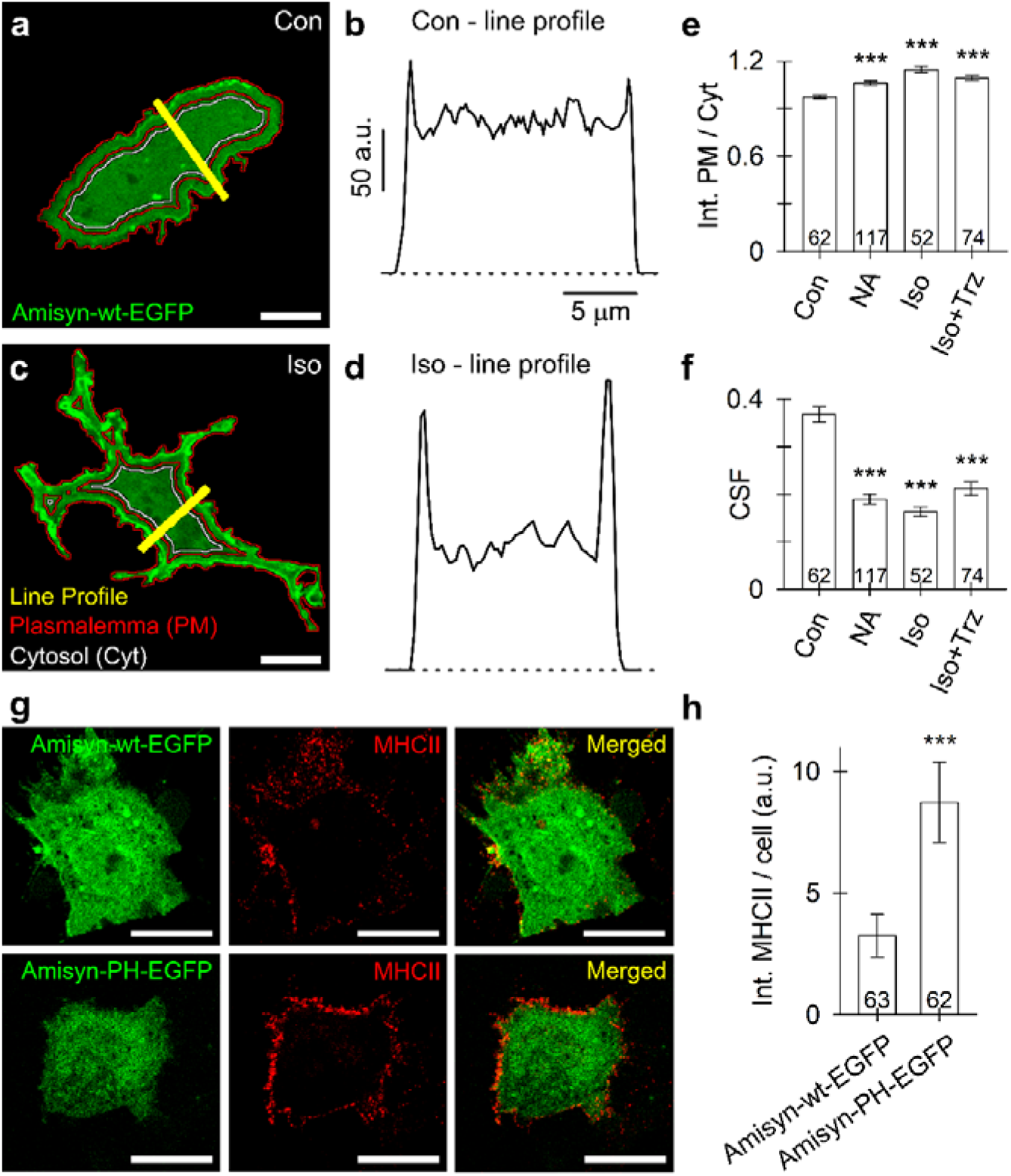
β-Adrenergic treatment favours astrocyte arborization and enhances the distribution of wild-type (wt) amisyn to the plasmalemma (PM), and overexpression of wt amisyn reduces interferon γ (IFNγ)-induced surface expression of MHCII compared with overexpression of the amisyn pleckstrin homology (PH) mutant. (**a, c**) Confocal micrographs depicting the subcellular distribution of amisyn-wt-enhanced green fluorescent protein (EGFP) in extracellular solution (ECS)-treated controls (Con; **a**) and isoprenaline (Iso)-treated astrocytes (**c**). Note the morphological change between the cell image in (**a**) and (**c**). Yellow lines indicate fluorescence intensity line profiles drawn across astrocytes. The red-outlined area indicates a 2-µm thick band used to measure amisyn-wt-EGFP fluorescence at the plasmalemma, whereas the white-circled region depicts fluorescence measured in the cytosol (cell interior). Scale bars: 10 µm. (**b, d**) Line profiles (drawn from **a** and **c**). Note the higher fluorescence intensity peaks in astrocytes treated with Iso compared with the ECS-treated control. Higher peaks indicate relatively more amisyn-wt-EGFP associated with the plasmalemma in Iso-treated astrocytes. (**e**) Ratio between amisyn-wt-EGFP fluorescence at the plasmalemma and in the cytosol in controls, and in astrocytes treated with NA, Iso or co-treated with Iso + Trz. Note an increase in the PM/Cyt ratio in astrocytes treated with (non-)selective agonists of β-adrenergic receptors. (**f**) The cell shape factor (CSF = 4 × π × *S*/*p*^2^, where *S* is the surface area and *p* is the cell perimeter) was calculated in ECS-treated controls and in astrocytes treated with NA, Iso or co-treated with Iso + Trz. A decrease in the CSF indicates an increase in arborized (less circular) morphology. \*\*\**P* < 0.001 (ANOVA on ranks followed by Dunn’s test; **e, f**). (**g**) Confocal micrographs of surface-expressed MHCII immunolabelled by the anti-MHCII antibody and by the corresponding fluorescent secondary antibody (MHCII, red) in astrocyte transfected to overexpress EGFP-tagged (green) amisyn-wt (top) or amisyn-PH (bottom). Cells were examined ∼48 h post transfection and IFNγ treatment. Scale bars (**g**): 10 µm. (**h**) Mean (± SEM) MHCII fluorescence per cell in astrocytes overexpressing amisyn-wt-EGFP or amisyn-PH-EGFP. Numbers at the bottom of the bars indicate the number of cell images analysed. \*\*\**P* < 0.001 (Mann-Whitney U test).

We then assessed the effect of amisyn on MHCII surface expression in IFNγ-treated astrocytes. Peripheral MHCII fluorescence was approximately threefold lower in cells overexpressing wt amisyn than in those expressing the PI(4,5)P_2_-binding-deficient PH mutant (3 ± 1 versus 9 ± 2 arbitrary units [a.u.]; *P* < 0.05; Fig. 4g, h), consistent with inhibition of exocytosis of MHCII-containing vesicles.

Because MHCII co-localizes with lysosomal markers, we also monitored CD63-pHuji-labelled vesicles. Super-resolution imaging confirmed MHCII and CD63 co-localization (Extended Data Fig. 5a–c). The plasmalemma-to-cytosol CD63-pHuji ratio was lower in astrocytes expressing wt amisyn than in those expressing the PH mutant (0.89 ± 0.02 versus 1.14 ± 0.02; *P* < 0.001; Extended Data Fig. 5d, e), indicating that amisyn suppresses lysosomal exocytosis and the surface delivery of both MHCII and CD63.

### Wild-type amisyn inhibits single-vesicle secretion in astrocytes

To assess the effect of amisyn on vesicle discharge, astrocytes expressing atrial natriuretic peptide (ANP) tagged with emerald green fluorescent protein (ANP.emd)^43^ were co-transfected with wt or PH-mutant amisyn (pAmisyn-wt-RFP or pAmisyn-PH-mCherry; Fig. 5a) and stimulated with ATP (100 µM, 4 min) to invoke Ca^2+^-dependent vesicle secretion. Secretory events appeared as abrupt fluorescence losses occurring within the 0.5-s sampling interval (Fig. 5b, c).

**Fig. 5.**
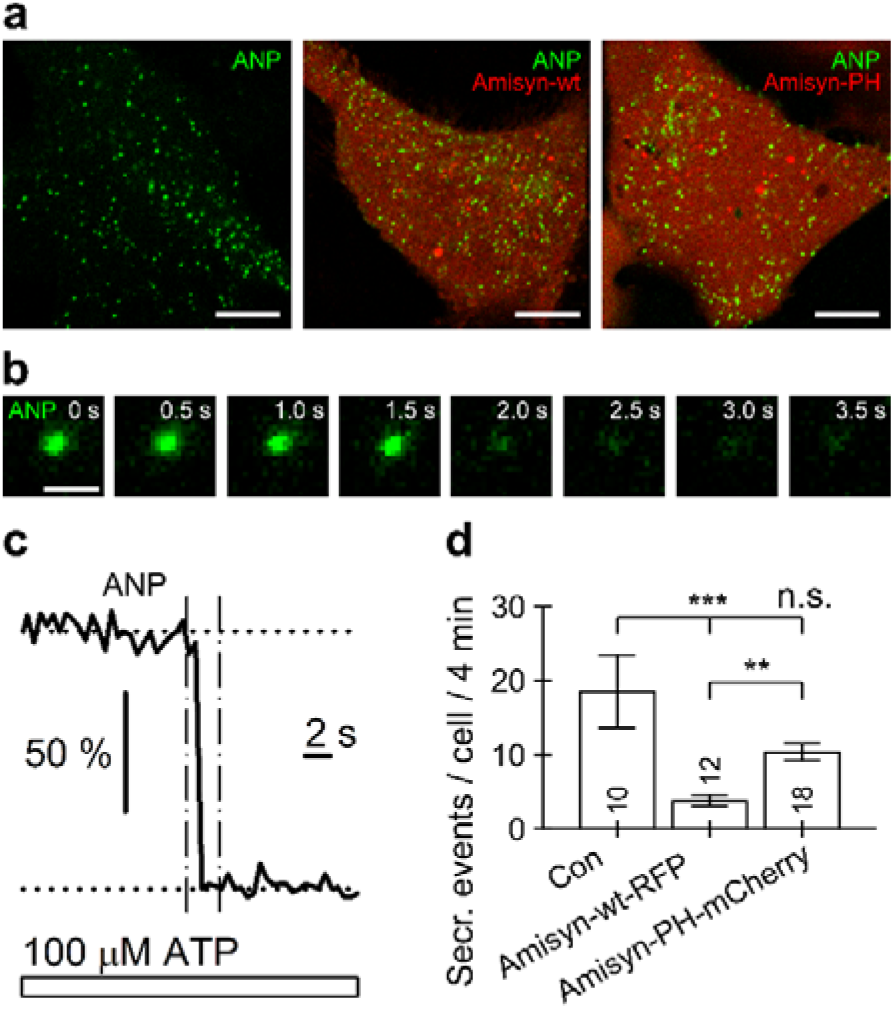
Overexpression of wild-type (wt) amisyn reduces the occurrence of ATP-evoked exocytotic secretory events from single vesicles in astrocytes. (**a**) Confocal images of live astrocytes transfected to (co-)express (left) ANP.emd (emerald GFP-tagged atrial natriuretic peptide; green), (middle) ANP.emd and Amisyn-wt-RFP, and (right) ANP.emd and Amisyn-PH-mCherry. (**b**) Sequence of confocal images depicting rapid (∼0.5 s) discharge of fluorescent cargo during cell stimulation with 100 μM ATP. Time marks in the top right corners indicate the time relative to the arbitrary starting point (0 s). Scale bars: 10 μm (**a**) and 1 μm (**b**). (**c**) Normalized time-dependent change in vesicle fluorescence (from **b**) acquired in the circular region of interest (r = 6 pixels, 3.3 μm). The horizontal dotted lines indicate the minimum and maximum fluorescence levels. The vertical dashed lines indicate the time period during which vesicle fluorescence was measured from the images displayed in (**b**). The addition of ATP is indicated by the horizontal rectangle. (**d**) The number (mean ± SEM) of ATP-evoked secretory events in individual cells within a 4-min window. Overexpression of wt amisyn reduced the number of single-vesicle secretory events observed. The numbers at the bottom of the bars indicate the number of astrocytes analysed. \*\**P* < 0.01, \*\*\**P* < 0.001 (ANOVA on ranks followed by Dunn’s test).

ATP-evoked events were significantly reduced by wt amisyn, from 18 ± 5 to 4 ± 1 events per cell per unit time, whereas the PH mutant had a weaker effect (11 ± 1; *P* < 0.01 and *P* < 0.001; Fig. 5d). Thus, wt, but not PH-mutant, amisyn inhibits ATP-evoked single-vesicle secretion in astrocytes.

### Amisyn is required for **β**-adrenergic regulation of the exocytotic fusion pore

We performed high-resolution cell-attached *C*_m_ measurements in 171 astrocytes from 26 independent cultures to determine how amisyn regulates single-vesicle exocytosis, resulting in 23.5 h of recordings. Cells overexpressed EGFP, wt amisyn, or PH-mutant amisyn, or were treated with amisyn small interfering RNA (siRNA), with or without Iso (Fig. 6a; Table 2). Amisyn knockdown was confirmed by reduced amisyn-EGFP fluorescence and a decrease in endogenous amisyn mRNA to 30.5% ± 3.0% of control levels (Extended Data Fig. 6).

**Fig. 6.**
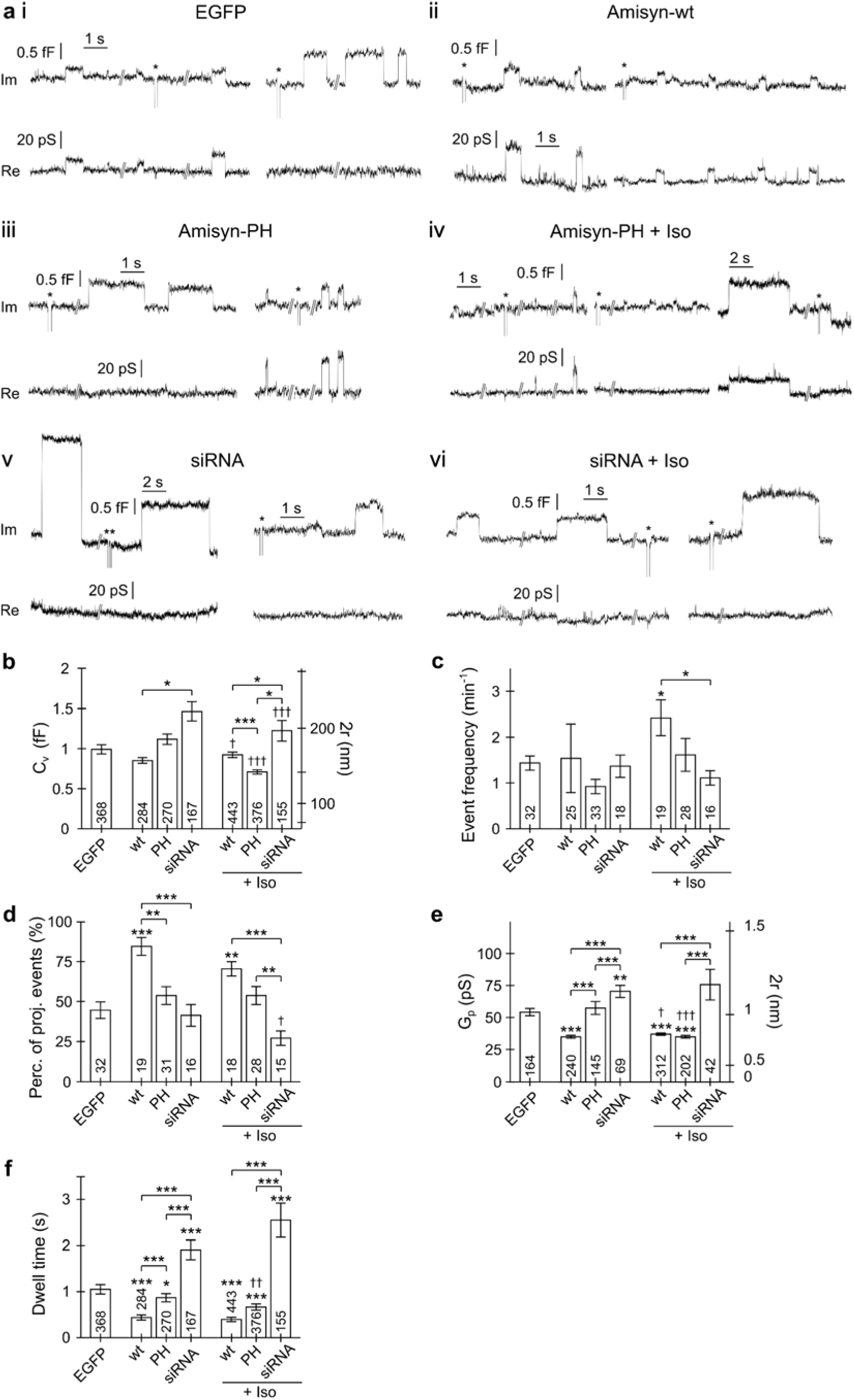
β-Adrenergic regulation of the exocytotic fusion pore depends on amisyn. (**a**) Representative recordings of transient vesicle exocytosis in enhanced green fluorescent protein (EGFP)-transfected controls (**i**), amisyn-wild-type (wt)-EGFP (Amisyn-wt, wt) transfected astrocytes (**ii**), amisyn-PH-EGFP (Amisyn-PH, PH) transfected astrocytes (**iii**), astrocytes treated with isoprenaline (Iso) (**iv**), and amisyn knockdown (small interfering RNA [siRNA]) astrocytes (**v**), treated with Iso (**vi**). Asterisks (*) denote calibration pulses. (**b–f**) Graphs depicting (mean ± SEM) the vesicle capacitance (*C*_v_; **b**), the event frequency (**c**), the percentage of Re-projected events (vesicles establishing a narrow fusion pore) (**d**), fusion-pore conductance (*G*_p_; **e**) and open pore dwell time (**f**). The graphs depicting vesicle capacitance (*C*_v_; **b**) and fusion-pore conductance (*G*_p_; **e**) display the corresponding vesicle diameter or pore diameter on the ordinate on the right. Numbers at the bottom of the bars denote the number of events (**b, e, f**) or recordings (**c, d**) analysed. The symbols above the bar plots (**b–f**) depict four sets of statistical comparisons: symbols on the top of the individual bars denote (*) comparisons versus the EGFP group used as an experimental control or (†) comparisons versus the analogous Iso-treated group (above 3 bars on the right). Two additional sets statistically compare the effects of differential amisyn expression (wt, PH, siRNA) in astrocytes not treated (symbols above 3 bars in the middle) or treated with Iso (symbols above 3 bars on the right). Pairwise comparisons (Dunn’s tests) were conducted after ANOVA on ranks and were Bonferroni corrected. */†*P* < 0.05, **/††*P* < 0.01, ***/†††*P* < 0.001.

**Table 2.** Membrane capacitance analysis of astrocytes overexpressing EGFP, amisyn-wt-EGFP (wt), amisyn-PH-EGFP (PH), or amisyn siRNA (siRNA), under basal conditions or following isoprenaline stimulation (+Iso). Analysis of elementary exocytotic activity of full (Full) and transient (Trans.) exocytotic (Exo) events. (Table 2a) Overview. (Table 2b) Descriptive statistics (mean ± SEM) of transient exocytotic events in astrocytes overexpressing EGFP, amisyn-wt-EGFP (wt), amisyn-PH-EGFP (PH), or amisyn siRNA (siRNA), under basal conditions or after isoprenaline stimulation (+Iso). (Table 2c) Pairwise statistical comparisons of transient exocytotic events in transfected cells.

| Table 2a | Number of cells | Effective time (min) | Total event count | Total event frequency (min<sup>-1</sup>) | Event count |  | Percentage (%) |  | Event frequency (min<sup>-1</sup>) |  |
| --- | --- | --- | --- | --- | --- | --- | --- | --- | --- | --- |
| Treatment |  |  |  |  | Full Exo | Trans. Exo | Full Exo | Trans. Exo | Full Exo | Trans. Exo |
| EGFP | 32 | 256.3 | 439 | 1.71 | 71 | 368 | 16.2 | 83.8 | 0.28 | 1.44 |
| wt | 25 | 184.2 | 315 | 1.71 | 31 | 284 | 9.8 | 90.2 | 0.17 | 1.54 |
| PH | 33 | 292.4 | 337 | 1.15 | 67 | 270 | 19.9 | 80.1 | 0.23 | 0.92 |
| siRNA | 18 | 122.2 | 220 | 1.80 | 53 | 167 | 24.1 | 75.9 | 0.43 | 1.37 |
| wt + Iso | 19 | 182.9 | 498 | 2.72 | 55 | 443 | 11.0 | 89.0 | 0.30 | 2.42 |
| PH + Iso | 28 | 232.9 | 463 | 1.99 | 87 | 376 | 18.8 | 81.2 | 0.37 | 1.61 |
| siRNA + Iso | 16 | 139.5 | 222 | 1.59 | 67 | 155 | 30.2 | 69.8 | 0.48 | 1.11 |

EGFP, enhanced green fluorescent protein; wt, wild-type; PH, homology; siRNA, small interfering RNA; Iso, isoprenaline.

| **Table 2b** | **Number of events** | **Number of cells** | **$C_v$ (fF)** | **Event frequency (events/min)** | **Percentage of projected events (%)** | **$G_p$ (pS)** | **$G_p/C_v$ (pS/fF)** | **Dwell time (s)** |
| --- | --- | --- | --- | --- | --- | --- | --- | --- |
| EGFP | 368 | 32 | $0.99 \pm 0.05$ | $1.43 \pm 0.15$ | $44.5 \pm 5.1^{***}$ | $54.2 \pm 2.8^{***}$ | $40 \pm 2.1^{***}$ | $1.05 \pm 0.1^{***}$ |
| wt | 284 | 25 | $0.85 \pm 0.03$ | $1.54 \pm 0.74$ | $84.5 \pm 5.5$ | $35 \pm 1.1$ | $41.6 \pm 0.6$ | $0.43 \pm 0.1$ |
| PH | 270 | 33 | $1.11 \pm 0.06$ | $0.92 \pm 0.15$ | $53.7 \pm 5.5^{**}$ | $57.4 \pm 5^{***}$ | $40.5 \pm 2.1^{***}$ | $0.86 \pm 0.1^{***}$ |
| siRNA | 167 | 18 | $1.46 \pm 0.11^*$ | $1.36 \pm 0.24$ | $41.3 \pm 6.8^{***}$ | $70.4 \pm 4.7^{***}$ | $42.9 \pm 2.9$ | $1.9 \pm 0.2^{***}$ |
| wt + Iso | 443 | 19 | $0.92 \pm 0.03$ | $2.42 \pm 0.38$ | $70.4 \pm 4.4$ | $37 \pm 0.8$ | $37.5 \pm 0.7^{***}$ | $0.39 \pm 0.04$ |
| PH + Iso | 376 | 28 | $0.7 \pm 0.02^{***}$ | $1.61 \pm 0.35$ | $53.7 \pm 5.6^{**}$ | $35 \pm 1$ | $37 \pm 1^{***}$ | $0.66 \pm 0.06$ |
| siRNA + Iso | 155 | 16 | $1.22 \pm 0.12$ | $1.11 \pm 0.15$ | $27 \pm 4.5^{***}$ | $75.7 \pm 11.9^{***}$ | $39.7 \pm 3.2$ | $2.54 \pm 0.37^{***}$ |

| Table 2c | $C_v$ (fF) | Event frequency<br>(events/min) | Percentage of<br>projected events<br>(%) | $G_p$ (pS) | Dwell time<br>(s) |
| --- | --- | --- | --- | --- | --- |
| Comparison |  |  |  |  |  |
| EGFP - wt | >0.5 | >0.5 | <0.001*** | <0.001*** | <0.001*** |
| EGFP - PH | 0.054 | 0.136 | >0.5 | 0.33 | 0.012 |
| EGFP - siRNA | <0.001*** | >0.5 | >0.5 | 0.003** | <0.001*** |
| EGFP - wt + Iso | 0.084 | 0.049* | 0.009** | <0.001*** | <0.001*** |
| EGFP - PH + Iso | 0.002* | >0.5 | >0.5 | <0.001*** | <0.001*** |
| EGFP - siRNA + Iso | >0.5 | >0.5 | 0.232 | 1 | <0.001*** |
| wt - wt + Iso | 0.019* | 0.346 | 0.058 | 0.048 | >0.5 |
| PH - PH + Iso | <0.001*** | 0.067 | >0.5 | <0.001*** | 0.002** |
| siRNA - siRNA + Iso | <0.001*** | 0.399 | 0.097 | 0.137 | >0.5 |
| wt - PH | >0.5 | >0.5 | 0.001** | <0.001*** | <0.001*** |
| wt - siRNA | 0.018* | >0.5 | <0.001*** | <0.001*** | <0.001*** |
| PH - siRNA | 0.204 | 0.347 | >0.5 | <0.001*** | <0.001*** |
| wt + Iso - PH + Iso | 0.036* | 0.424 | 0.118 | <0.001*** | <0.001*** |
| wt + Iso - siRNA + Iso | 0.018* | 0.017* | <0.001*** | <0.001*** | <0.001*** |
| PH + Iso - siRNA + Iso | 0.039* | >0.5 | 0.008** | <0.001*** | <0.001*** |

$C_v$ , vesicle capacitance; $G_p$ , pore conductance; EGFP, enhanced green fluorescent protein; wt, wild-type; PH, pleckstrin homology; siRNA, small interfering RNA; Iso, isoprenaline. Bonferroni-corrected $P$ values result from individual comparisons (Dunn's test, t test) after ANOVA on ranks or one-way ANOVA. \* $P$ < 0.05; \*\* $P$ < 0.01; \*\*\* $P$ < 0.001.

Although mean vesicle capacitance *C*_v_ did not differ significantly from EGFP controls, wt amisyn reduced the contribution of larger vesicles relative to amisyn knockdown, both with and without Iso (Fig. 6b; Extended Data Fig. 7). The relative frequency of full-fusion events was higher after amisyn knockdown than in cells overexpressing wt amisyn, indicating that amisyn restrains progression from transient to complete vesicle fusion (Extended Data Fig. 8). Notably, Iso only slightly, but statistically significantly, reduced vesicle size after amisyn knockdown. This effect was not due to altered β_1_-adrenergic receptor expression in siRNA amisyn-treated cells, which remained unchanged (Extended Data Fig. 6a).

Amisyn expression did not significantly alter the overall frequency of transient events (Fig. 6c). However, Iso increased their relative frequency in cells overexpressing wt amisyn, consistent with repeated opening and closing of less-stable fusion pores.

Fusion-pore analysis identified amisyn as a key regulator of fusion-pore electrical geometry and kinetics (Fig. 6d–f). Wild-type amisyn increased the proportion of narrow-pore events (transient fusions exhibiting a projection between Im and Re admittance signals), reduced fusion-pore conductance (*G*_p_), and shortened pore dwell time, whereas amisyn knockdown widened the pore, prolonged dwell time, and promoted full fusion (Extended Data Fig.8). Iso reduced fusion-pore conductance in cells expressing PH-mutant amisyn, presumably via endogenous amisyn, but had no additional effect in cells overexpressing wt amisyn or in cells lacking amisyn.

These findings show that amisyn is required for β-adrenergic control of the exocytotic fusion pore. Wild-type amisyn suppresses fusion of larger vesicles, narrows the pore, shortens its open time, and limits full fusion, whereas amisyn depletion produces the opposite effects.

## Discussion

Loss of noradrenergic tone due to locus coeruleus degeneration is increasingly recognized as a driver of astrocyte-mediated neuroinflammation in neurodegenerative disorders. Here, we show that β-adrenergic stimulation requires amisyn to reduce MHCII surface expression in reactive astrocytes by inhibiting exocytosis of large lysosome-like vesicles, decreasing fusion-pore conductance and dwell time, and promoting amisyn localization at the plasma membrane.

Reactive astrocytes undergo extensive morphological, molecular, and functional changes in response to pathological insults^8^, including acquisition of a pro-inflammatory phenotype characterized by increased expression of the antigen-presenting molecule MHCII^35^. We therefore investigated mechanisms that suppress MHCII surface expression in IFNγ-treated astrocytes, a model previously examined using vesicle-dynamics analysis^35^, immunoelectron microscopy^44^, live-cell immunocytochemistry, and electrophysiology^18^.

NA, an endogenous neurotransmitter with anti-inflammatory properties^45^, inhibited IFNγ-induced MHCII surface expression in human and rat astrocytes (Fig. 1), consistent with earlier findings^36^. Pharmacological experiments using selective agonists and antagonists showed that this effect was mediated by β- but not α-adrenergic receptors (Fig. 2). The involvement of β-adrenergic signalling was further supported by the inhibitory effect of dbcAMP, indicating that increased intracellular cAMP contributes to reduced MHCII surface expression (Fig. 2). In vivo, this pathway may limit astrocytic antigen presentation and thereby counteract neuroinflammation associated with neurodegeneration^3^.

cAMP may regulate MHCII surface expression through both transcriptional and vesicular mechanisms. At the transcriptional level, the inducible cAMP early repressor inhibits cAMP response element-binding protein activity and thereby interferes with the class II transactivator CIITA, a major regulator of MHCII expression^17,46^. In parallel, cAMP-dependent changes in vesicle trafficking may reduce the delivery of MHCII-containing organelles to the plasma membrane.

Astrocytic lysosomes undergo regulated exocytosis, releasing low-molecular-weight signalling molecules^47–49^ and transporting MHCII to the cell surface^18^. Reduced vesicle delivery contributes to lower surface expression in astrocytes^35^.

Depending on the stimulus, astrocytic lysosomes undergo either transient or full-fusion exocytosis^47,50^, with full fusion primarily supporting plasma membrane repair^50,51^. Unlike neurons, astrocytes spontaneously internalize fluorescent styryl dyes, rapidly labelling vesicles positive for sialin, CD63/LAMP3, and VAMP7, markers of late endosomes and lysosomes. These findings indicate that a subset of astrocytic lysosomes rapidly recycles after Ca^2+^-dependent exocytosis^52^.

Even in unstimulated astrocytes, spontaneous microdomain-localized increases in intracellular Ca^2+^ occur at a frequency of 0.30 ± 0.02 events/min^53,54^. These Ca^2+^ signals may trigger lysosomal exocytosis, which depends on a lysosomal-specific set of SNARE proteins^55^.

Consistent with previous membrane capacitance studies^39,56–59^, we recorded both transient and full-fusion exocytotic events; transient events predominated in astrocytes (Fig. 3), as reported previously^59,60^. In IFNγ-treated astrocytes, transiently fusing vesicles had a larger estimated diameter, reflected by increased vesicle capacitance (*C*_v_), indicating a shift towards larger exocytotic compartments (Fig. 3). Their mean diameter of 173 ± 6 nm was consistent with LAMP1-positive lysosomal compartments^60^, supporting the role of lysosome-like vesicles in MHCII trafficking to the plasma membrane^18,61^.

β-Adrenergic co-treatment with NA or Iso reduced the frequency of fusion events involving larger vesicles (Fig. 3), suggesting selective inhibition of lysosome-like vesicle engagement with the plasma membrane. This finding is consistent with reports that increased intracellular cAMP favours transient fusion of smaller vesicles in astrocytes and pituitary cells^62,63^. Thus, β-adrenergic signalling appears to reshape the vesicle population undergoing exocytosis in IFNγ-reactive astrocytes by restricting the access of larger lysosome-like compartments to the plasma membrane.

Amisyn is a likely mediator of this β-adrenergic inhibition of MHCII presentation through its regulation of the exocytotic fusion pore^64^. Our results show for the first time that amisyn controls both fusion-pore width and opening duration, as assessed by fusion-pore conductance (*G*_p_) and dwell time, respectively (Fig. 6). Increased intracellular cAMP has previously been shown to recruit amisyn to vesicle fusion sites^41^, where it may regulate transient (kiss-and-run) exocytosis in pancreatic β-cells and neuroendocrine cells ^62,65^.

Increased cAMP, induced by forskolin or β-adrenergic agonists, inhibits phosphoinositide turnover and intracellular Ca² mobilization^66,67^. cAMP may stabilize PI(4,5)P_2_ at exocytotic sites^68^, promoting the recruitment of amisyn through its PH domain. Alternatively, cAMP may act through Epac2, which translocates to the plasma membrane and interacts with the secretory machinery^69^. In β-cells, Epac2 recruits amisyn and dynamin-1, thereby restricting fusion pore expansion^41^. Unlike the amisyn-related protein tomosyn (STXBP5)^70,71^, amisyn does not exhibit a protein kinase A phosphorylation site, according to analysis of its amino-acid sequence using NetPhos (https://services.healthtech.dtu.dk/services/NetPhos-3.1/ accessed on 16 April 2024).

We further show that amisyn is enriched at the astrocyte plasma membrane (Fig. 4; Extended Data Fig. 5), as reported in other cell types^25,28^. At this site, it may increase the abundance of fusion-inactive SNARE complexes, thereby suppressing vesicle discharge^28^. Although amisyn lacks a transmembrane domain and recognizable lipidation motifs, its intact PH domain acts as a PI(4,5)P_2_-dependent effector that enables plasma membrane association and is required for its inhibitory effect on vesicle fusion (Extended Data Fig. 5)^28,72–74^.

In PC12 and chromaffin cells, expression of full-length amisyn reduced the number of exocytotic events via syntaxin-1-dependent and -independent mechanisms^27^. In chromaffin cells, expression of wt amisyn inhibited basal and stimulated exocytosis. Similarly, in astrocytes, overexpression of amisyn-wt-EGFP, but not amisyn-PH-EGFP, inhibited exocytosis, as indirectly indicated by the reduced surface expression of MHCII (Fig. 4) and CD63-pHuji (Extended Data Fig. 5), and directly by the reduced occurrence of unitary secretory vesicle discharge events (Fig. 5).

Beyond reducing exocytotic activity, amisyn strongly regulates fusion-pore behaviour. In chromaffin cells, amisyn prolongs the amperometric foot signal preceding transmitter release, suggesting delayed fusion-pore expansion and persistence of a narrow-conductance intermediate^27,75^. In astrocytes, amisyn overexpression increased the proportion of transient exocytotic events, whereas amisyn knockdown increased *G*_p_, prolonged pore dwell time, and promoted progression to full fusion (Fig. 6). These findings indicate that amisyn suppresses exocytosis by limiting fusion-pore expansion and restraining the transition from transient to full-fusion exocytosis.

Fusion-pore width is likely regulated by the number of fusogenic trans-SNARE complexes assembled between the vesicle and plasma membrane. A small number of SNARE complexes supports the formation of narrow, transient pores, whereas greater SNARE assembly promotes pore expansion and full vesicle collapse^76^. Amisyn may therefore restrict SNARE complex assembly, producing narrower fusion pores, as indicated by lower fusion-pore conductance (*G*_p_) and higher frequency of narrow fusion pores (Fig. 6).

At rest, larger vesicles generally form wider and more stable fusion pores than smaller vesicles and are therefore more likely to progress to full fusion (Extended Data Fig. 3c; Fig. 3)^40,60,77^. By limiting productive SNARE assembly, amisyn may reduce pore dilation and promote narrow, transient fusion pores, thereby preventing complete vesicle collapse.

Our findings support a concentration-dependent inhibitory action of amisyn on SNARE-mediated fusion. Like tomosyn, syntaphilin, and dominant-negative SNARE constructs^60,71,78^, amisyn may function as a decoy SNARE protein^79^. Moderate amisyn levels may constrain fusion-pore expansion, whereas higher levels may suppress fusion altogether by preventing the formation of sufficient fusogenic SNARE complexes. The similarity between astrocytes overexpressing amisyn and those expressing dominant-negative SNARE peptides further supports a role for amisyn in stabilizing narrow, release-limiting pores, particularly in large vesicles^60^.

Fusion-pore constriction is not exclusively restricted to protein-mediated mechanisms. Changes in membrane lipid composition, particularly cholesterol enrichment, also reduce fusion-pore conductance and oppose pore expansion. In astrocytes and lactotrophs, vesicular cholesterol exerts a constrictive effect on the fusion pore, whereas cholesterol depletion promotes pore widening and secretion^80^. This effect is thought to arise from membrane-mechanical forces at the vesicle–plasma membrane interface that stabilize narrow-pore configurations.

Consistent with a SNARE-based mechanism, amisyn also reduced the probability of full fusion (Extended Data Fig. 8). By limiting productive SNARE complex assembly and pore expansion, amisyn shifted exocytosis away from complete vesicle collapse towards transient fusion. Conversely, amisyn depletion promoted full fusion, indicating that amisyn regulates not only fusion-pore geometry and kinetics but also the transition between reversible and irreversible exocytosis. Thus, although acting through distinct molecular pathways, amisyn and cholesterol converge on a common outcome: stabilization of narrow fusion pores and restriction of full-fusion exocytosis.

In conclusion, β-adrenergic signalling targets amisyn, a negative regulator of astrocyte exocytosis, by limiting fusion efficiency and fusion-pore expansion. Increased intracellular cAMP may promote amisyn localization near MHCII-positive vesicles preparing for exocytosis, thereby reducing MHCII delivery to the plasma membrane and antigen presentation. This β-adrenergic pathway may protect against astrocyte-mediated neuroinflammation, whereas loss of noradrenergic signalling after locus coeruleus degeneration may weaken this restraint in neurodegenerative diseases.

## Abbreviations

ANP: atrial natriuretic peptide
a.u.: arbitrary units (intensity)
BSA: bovine serum albumin
CD63: cluster of differentiation protein 63 (LAMP-3, TSPAN30)
CIITA: the MHCII transactivator
*C*_m_: membrane capacitance
CNS: central nervous system
CSF: cell shape factor
*C*_v_: vesicle capacitance
DAPI: 4′,6-diamidino-2-phenylindole fluorescent stain, marking nuclei
dbcAMP: dibutyryl cyclic adenosine monophosphate, membrane-permeable analogue of cAMP
DIC: differential interference contrast
DPSS: diode-pumped solid-state
ECS: extracellular solution
EGFP: enhanced green fluorescent protein
*G*_p_: pore conductance
IFNγ: interferon γ
Im: imaginary part of the admittance signal, proportional to *C*_m_
Iso: isoprenaline, a β-adrenergic agonist
JAK: Janus kinase
MHCII: major histocompatibility complex II
NA: noradrenaline
PBS: phosphate-buffered saline
PC12: pheochromocytoma-derived cell line of the rat adrenal medulla
PE: phenylephrine, an α-adrenergic agonist
PH: pleckstrin homology
PI(4,5)P_2_: phosphatidylinositol 4,5-bisphosphonate
Pro: propranolol, a β-adrenergic antagonist
Re: real part of the admittance signal
RMS: root means square (signal magnitude)
RT: room temperature (∼22°C)
SIM: structured illumination microscopy
siRNA: small interfering RNA
STXBP6: syntaxin-binding protein 6
Trz: terazosin, an α-adrenergic antagonist
wt: wild-type.

## Methods

### Culture of primary rat astrocytes

Primary astrocyte cultures were prepared from cerebral cortices of 2- to 3-day-old female Wistar rats as described previously^82^. Animal handling complied with European and Slovenian legislation (Official Gazette of the RS 38/13 and official consolidated text 21/18, 92/20, 159/21; UVHVVR, permit no. U34401-27/2025/4, U34401-26/2025/).

Isolated cells were maintained in high-glucose Dulbecco’s modified Eagle’s medium, supplemented with 10% foetal bovine serum, 1 mM sodium pyruvate, 2 mM L-glutamine, and 25 µg/ml penicillin/streptomycin (culture medium) in an atmosphere of 5% CO_2_/humidified air (95%) at 37°C. Subconfluent cultures were shaken at 225 rotations/min overnight, followed by a medium exchange. This was repeated three times. After enrichment, astrocytes were detached from the culture flask using 0.1% trypsin and 0.04% EDTA in Hank’s balanced salt solution and subcultured into culture tubes. The culture medium was exchanged 3 times/week. One to four days before the experiments/measurements, astrocytes were plated on individual 22-mm diameter poly-L-lysine-coated glass coverslips at varying densities, depending on the type of experiment.

To study the effects of adrenergic signalling on IFNγ-induced MHCII surface expression, we (co-)incubated astrocytes for 48 h at 37°C with IFNγ (60 U/ml or 100 U/ml; U-Cytech, Utrecht, the Netherlands) and diverse adrenergic agents. We used 100 μM phenylephrine hydrochloride (PE) and 2 μM propranolol hydrochloride (Pro) for selective α-adrenergic stimulation, 100 μM isoprenaline hydrochloride (Iso) and 2 μM terazosin (Trz) for selective β-adrenergic stimulation and either 10 μM or 100 μM noradrenaline hydrochloride for nonselective, α- and β-adrenergic stimulation. All chemicals were purchased from Merck (Darmstadt, Germany) unless stated otherwise.

### Culture of primary human astrocytes

A cryopreserved vial of human cortical GFAP-positive astrocytes (10^6^ cells) was purchased from ScienCell (Carlsbad, CA, USA; cat. no. 1800) via Provitro (Berlin, Germany). The cell culture originated from a healthy male donor in the 20th week of gestation (lot. no. 41331; P1, cryopreserved on 13 December 2024). The culture was thawed, propagated and maintained according to the manufacturer’s instructions using proprietary Astrocyte Medium (ScienCell, cat. no. 1801). MHCII expression was induced by 100 U/ml of *Escherichia coli*-derived human IFNγ (cat. no. HC2030-01, Hycult Biotech, Uden, the Netherlands).

### Solutions

The extracellular solution (ECS) for live-cell imaging and electrophysiological measurements consisted of 130 mM NaCl, 5 mM KCl, 2 mM CaCl_2_, 1 mM MgCl_2_, 10 mM D-glucose, and 10 mM HEPES/NaOH (pH 7.2). Osmolarity (300 ± 15 mOsm) was measured with a freezing-point osmometer (Osmomat 030, Gonotec, Berlin, Germany). For visualization of CD63-pHuji in acidified compartments^83^, we substituted a fraction of the NaCl with NH_4_Cl in the ECS (80 mM NaCl + 50 mM NH_4_Cl). This solution was added to the ECS at a 1:4 dilution (to achieve 10 mM NH_4_Cl contact concentration) just before beginning the measurements.

### Plasmids and cell transfection

To study the subcellular distribution of amisyn and its involvement in the regulation of exo-and endocytosis, we transfected astrocytes with plasmids encoding enhanced green fluorescent protein (EGFP)-tagged wild-type (wt) amisyn and amisyn with a mutated pleckstrin-homology (PH) domain^28^. To study how overexpressed amisyn variants (wt and PH) affect the occurrence of secretory events, we (co-)transfected astrocytes with plasmids encoding atrial natriuretic peptide C-terminally tagged with emerald green fluorescent protein (ANP.emd; a gift from Dr Ed Levitan, University of Pittsburgh, Pittsburgh, PA, USA)^43,84^, red fluorescent protein (RFP)-tagged wt amisyn (amisyn-wt-RFP) and mCherry-tagged amisyn with a mutated PH domain (amisyn-PH-mCherry). We prepared amisyn-PH-mCherry plasmid by exchanging the C-terminal EGFP tag on amisyn-PH-EGFP with the C-terminal mCherry tag. To optophysiologically monitor exocytotic surface deposition of lysosomal proteins, we co-transfected astrocytes with plasmids encoding CD63-pHuji^83^ and EGFP-tagged amisyn variants (wt and PH).

Astrocytes were co-transfected with the aforementioned plasmids using FuGENE 6 (Promega Corporation, Madison, WI, USA). Briefly, for each coverslip, 3 µl of FuGENE 6 was diluted in 100 µl of culture medium, mixed, and incubated for 5 min at room temperature (RT). Then 0.8 µg of DNA was added (at a ratio of 1:3.75; DNA: FuGENE 6), mixed and incubated for a further 15 min at RT. Astrocytes were washed and incubated in 900 µl of fresh culture medium to which 100 µl of the transfection mixture was added. Transfected astrocytes were incubated for 24 h at 37°C in an atmosphere of 5% CO_2_/95% air. The transfection medium was replaced with fresh culture medium the next day, and transfected cells were observed 24–48 h after initiation of transfection.

We used a pulse transfection protocol (3–4 h of cell exposure to the transfection mixture) to mitigate amisyn expression and examined transfected cells earlier (16–24 h post-transfection time). We adopted a similar pulse transfection protocol to limit the negative influence of IFNγ treatment on the expression of plasmid-encoded proteins^85^ and started IFNγ treatment after a (3–4 h) pulse application of the transfection mixture to the cells. For co-transfection (transfection with two plasmids), we mixed half the amount of each plasmid (0.4 µg + 0.4 µg/coverslip) before exposure to the transfection reagent to promote protein co-expression in the same cells.

### Knockdown of amisyn via siRNA transfection

To estimate the time window necessary for amisyn knockdown, astrocytes were co-transfected with pre-designed siRNA targeting amisyn mRNA (rat STXBP6, Thermo Fisher Scientific, s169661) and pAmisyn-wt-EGFP. X-tremeGENE 360 Transfection Reagent (Merck), siRNA (20 nM; 20 pmol / coverslip) and pAmisyn-wt-EGFP (0.4 µg/coverslip) were diluted in 100 µl of serum-free culture medium, incubated for 20 min, and then applied to cells in serum-free culture medium at a final concentration of 20 nM siRNA. The siRNAs used included the target sequence: AAGGCGAAUAUUUAACUUATT). Cells were incubated for 4 or 12 h after co-transfection to allow knockdown to proceed. The transfection medium was then replaced with serum-containing medium, and cells were incubated for an additional 4–12 h before imaging or electrophysiological measurements.

### Quantitative real-time polymerase chain reaction

Total RNA was extracted from rat cortical astrocyte cell cultures using the E.Z.N.A. Total RNA Kit I (Omega Bio-Tek, Norcross, GA, USA). Total RNA was reverse transcribed into cDNA using the High-Capacity cDNA Reverse Transcription Kit (Thermo Fisher Scientific), according to the manufacturer’s instructions. Quantitative real-time polymerase chain reaction was performed using TaqMan Universal PCR Master Mix II (Thermo Fisher Scientific) and TaqMan Assays (Thermo Fisher Scientific): Stxbp6 (Amisyn; Rn01769493_m1), ADRB1 (Rn00824536_s1), and β-actin (Rn00667869_m1), according to the manufacturer’s instructions, in a QuantStudio 3 System (Thermo Fisher Scientific). Expression of target genes was normalized to expression of β-actin according to the equation target/reference = (1+EFF_reference_)^Ct_reference_)/(1+EFF_target_)^Ct_target_), where Ct is the quantification cycle and EFF is the amplification efficiency (expressed as a value between 0 and 1). EFF was determined with LinRegPCR software^86^.

### Immunocytochemistry

Immunocytochemical staining was performed as described previously^18^. PBS was used to wash the coverslips between individual steps, ∼3 min/wash. Blocking buffer and primary and secondary antibodies were diluted in a solution of 3% (w/v) BSA in PBS. With the exception of antibody incubation and blocking buffer, the immunolabelling procedure was performed at RT. First, astrocyte-loaded coverslips were washed once in PBS, fixed in formaldehyde (4% in PBS) for 15 min and permeabilized with 0.1% Triton X-100 for 10 min. Then astrocytes were washed four times and incubated with blocking buffer (10% goat serum, 1 h at 37°C) to reduce non-specific background staining. After an additional wash, primary antibodies were applied (overnight at 4°C). The next day, cells were washed four times before application of secondary antibodies (45 min at 37°C). After an additional four washes, we mounted the coverslips onto glass slides using SlowFade Gold antifade mountant with or without 4′,6-diamidino-2-phenylindole (DAPI; Thermo Fisher Scientific).

Depending on the primary antibody, we used goat anti-rabbit or anti-mouse secondary antibodies conjugated either to Alexa Fluor 546 or 488 (Thermo Fisher Scientific) at a dilution of 1:600. For quantification and characterization of MHCII-immunopositive vesicles, we used mouse monoclonal anti-MHCII (MRC-OX6; 1:100; ab23990, Abcam, Cambridge, UK) for rat astrocytes and mouse monoclonal anti-human HLA-DR antibodies (1:400, cat. no. 307602, Biolegend, San Diego, CA, USA) for human astrocytes. We used rabbit anti-Amisyn (Amichen #172; 1:500) to evaluate the expression of amisyn.

We conducted modified immunolabelling of live cells^18,87^ with a primary antibody (MRC-OX6) that binds to the extracellular domain of surface-exposed MHCII to visualize surface-expressed MHCII^18^. All labelling steps before fixation were performed on an ice-chilled surface with ice-cold solutions to prevent MHCII endocytosis. We used 3% BSA in PBS for washing; each wash lasted ∼3 min. Chilled coverslips were first washed once, incubated with 10% goat serum (∼3 min), washed again and incubated with primary antibodies (1:100; 15 min). Then coverslips were washed 3 times and incubated with secondary antibodies (1:600; 15 min). The coverslips were then washed 3 times, and cells were either supplied with ECS, transferred to the recording chamber and imaged, or washed again with PBS and fixed. Care was taken to avoid fast temperature changes to prevent cell blebbing^88^ and fixation was performed with 2% paraformaldehyde for 10 min. Coverslips were then washed three times with PBS and sealed to glass slides using SlowFade Gold antifade mountant with DAPI (Thermo Fisher Scientific).

### Image acquisition

#### Confocal microscopy

Fluorescently labelled cells were observed with a confocal microscope (LSM 780, Zeiss) using a plan apochromatic oil-immersion objective 63×/NA 1.4. Confocal images (single planes or z stacks) were acquired with a 405-nm diode-pumped solid-state (DPSS) laser, a 488-nm argon laser and a 561-nm DPSS laser, and the fluorescence emission was band-pass filtered at 440–480 nm (DAPI), 495–530 nm (amisyn-wt-EGFP or amisyn-PH-EGFP) and 565–615 nm (Alexa Fluor 546, CD63-pHuji), respectively.

#### Structured illumination microscopy

CD63-pHuji-expressing astrocytes containing immunofluorescent MHCII vesicles were transferred to the super-resolution Elyra PS.1 fluorescence microscope (Zeiss) and observed with an α-plan-apochromat oil-immersion differential interference contrast (DIC) objective (63×/1.40 Oil DIC M27). Immunofluorescent MHCII was excited by the 488 nm DPSS laser line, and emission was band-pass filtered at 495–575 nm. CD63-pHuji was excited by the 561-nm DPSS laser line, and emission was band-pass filtered at 570–650 nm. Emitted fluorescence was directed to the EMCCD Andor iXon 885 camera (Andor Technology, Belfast, UK), an electron-multiplying charge-coupled device, a type of digital camera sensor, to acquire 16-bit z stacks using five grating frequencies for structured illumination microscopy (SIM). The vertical (z) distance between the successive images in the z stack was 0.5 µm.

### Image analysis

Image analysis was performed using Fiji/ImageJ^89^ and MATLAB (MathWorks, Natick, MA, USA).

#### Quantification of MHCII expression in fixed, permeabilized astrocytes

MHCII expression was quantified as integrated density, thresholded at 30% of the intensity range (77 a.u.), divided by the number of astrocytes (DAPI-stained nuclei) in each image, as described previously^18^. We normalized the measured intensity to the mean intensity of individual IFNγ-treated groups (positive controls) to diminish batch-effect variance between different cell cultures and repetitions of experiments.

#### Quantification of MHCII surface expression

MHCII surface expression was quantified as peripheral MHCII immunofluorescence in individual astrocytes. A 2-μm band centred on the plasmalemma was manually outlined either in the confocal image (MHCII; if sufficient labelling was achieved) or in the DIC image, and MHCII fluorescence was calculated as the background-subtracted mean grey intensity within the band. This parameter was also used to discriminate between MHCII-positive and - negative astrocytes. The fluorescence threshold for the MHCII-positive count was selected on the basis of data obtained from a population of non-treated controls and IFNγ-treated astrocytes, so that almost no controls were MHCII-positive in contrast to 80% of IFNγ-treated astrocytes, consistent with the data from previous studies^35,36,90,91^.

#### Quantification of subcellular amisyn distribution

Subcellular amisyn distribution was examined in live amisyn-wt-EGFP-transfected cells (24 h) that were pre- and co-incubated with adrenergic agents. For the image analysis, we first acquired masks of individual cells by segmentation based on amisyn-EGFP (wt or PH) fluorescence, which indicated cytosolic amisyn distribution. For segmentation, we used the Huang auto-threshold^92^ in conjunction with basic morphological operations (open, close, fill holes) and a minimum size criterion for cell (particle) detection. We then visually examined the binary masks and manually corrected them if necessary. As a heuristic, we defined the 2-μm band from the surface to the cell interior as the plasmalemmal area (PM) and the subsequent 2-μm eroded mask image below the surface as the interior cytoplasmic area (CYT). Plasmalemmal association of amisyn was quantified as the ratio of mean fluorescence intensity within the two areas (*I*_PM_/*I*_CYT_), thresholded at 30 a.u. to limit the effects of background pixels due to imprecise segmentation.

The same mask images were used to analyse morphology, quantified by circularity or the CSF calculated as 4 × π × *S*/*p*^2^, where *S* is the surface area and *p* is the cell perimeter^42^. To exclude the effects of fine-scale segmentation on the estimation of the cell perimeter originating from differences in the subcellular distribution of EGFP-tagged amisyn variants (wt and PH mutant) and different expression (intensity levels), we used moderate morphological closing options in ImageJ (iterations = 5, count = 3).

#### Analysis of fluorescence co-localization

CD63 was validated as a suitable proxy for MHCII compartments by a pixel-based co-localization analysis as described previously^93^. For this, we analysed SIM images acquired with varying illumination and exposure times and applied an automatic thresholding method^94^. The resulting image masks were used to calculate fluorescence co-localization. Co-localization between immunolabelled proteins was expressed as the ratio of the co-localized pixel count to the total pixel count for a specific protein marker.

#### Lysosomal exocytosis estimated by subcellular distribution of CD63-pHuji fluorescence

We monitored the fluorescence of the tetraspanin CD63 (LAMP3) tagged with pHuji to quantify the inhibitory effect of overexpressed amisyn on endosomal/lysosomal exocytosis. CD63 is a suitable proxy because it co-localizes with MHCII-positive compartments, displays surface expression and is considered a marker of late endosomes/lysosomes or multivesicular bodies^95^. At low pH, the fluorophore pHuji is dim^96^ and not completely visualized inside the acidified compartments. Thus, we measured CD63-pHuji fluorescence either in fixed and permeabilized astrocytes or in live astrocytes bathed in 10 mM NH_4_Cl extracellular solution, which neutralized acidified vesicles and ensured reliable visualization of CD63-pHuji at the plasmalemma and inside vesicles^83^. Because the amount of protein expressed in individual transfected cells is stochastic, we used the ratio between the mean CD63-pHuji intensities (thresholded at 20 a.u. to limit the effects of background pixels due to imprecise segmentation) at the plasmalemma (within a 2 µm peripheral band) and inside the cell to estimate the relative fraction of membrane (surface) localization.

### Electrophysiology

Astrocyte-loaded coverslips were placed in the recording chamber, supplied with 200 μl of ECS and mounted on an inverted microscope (Zeiss Axio Observer, Zeiss). All experiments were performed at RT and measurements were obtained from non-stimulated recording periods (excluding non-acute adrenergic stimulation).

The exocytotic activity in individual astrocytes was recorded by the compensated cell-attached patch-clamp technique, enabling measurements of discrete stepwise increases and decreases in membrane capacitance (*C*_m_)^21,97^. We used standard-walled borosilicate glass pipettes (30-0058, Harvard Apparatus, Holliston, MA, USA), with a resistance of 2.5–3.5 MΩ, fire-polished and coated with the silicon resin Sylgard 184 (Dow Corning, Midland, MI, USA). The cell membrane was voltage-clamped by the dual-phase lock-in patch-clamp amplifier (SWAM IIC, Celica Biomedical, Slovenia) with a sine wave (0 mV ± 111 mV root mean square [RMS]) at a frequency (*f*) of 6400 Hz, which corresponds to the angular frequency (ω□=□2π*f*) of approximately 40,000 s^−1^. The phase angle of the lock-in amplifier was adjusted to nullify changes in the real (Re) part of the admittance signal in response to 10 fF calibration pulses in the imaginary (Im) part of the signal, triggered by a built-in calibration circuit to ensure correct phase angle settings^20^; continuously (150 ms square wave at 50% duty cycle) at the start of the recording and manually every□∼15 s. The signals were digitized at a sampling rate of 200 Hz. In accordance with good practice, all 3 signal traces (membrane current, Re, Im) were low-pass filtered at the Nyquist frequency: 100 Hz (4-pole Bessel filter, −3 dB).

In exocytotic events observed in Im (with or without a projection to Re), we calculated the vesicle capacitance using the equation: *C*_v_□=□[(Re^2^□+□Im^2^)/Im]/ω. The projection to Re indicated the formation of a narrow fusion pore that introduces a new resistive element into the electrical circuit^38,81,98^. *C*_m_ is proportional to the plasmalemmal area, the vesicle surface area and thus its diameter (*d*), therefore *C*_v_ can be determined from the equation *C*_v_□=□C_spec_π*d*^2^, where *C*_spec_ denotes the specific membrane capacitance. We calculated the vesicle diameter by assuming spherical geometry and by using a *C*_spec_ of 10 fF/µm^2^.^99^ The occurrence of Re projections enabled us to calculate the conductance of narrow fusion pores using the equation: *G*_p_□=□(Re^2^□+□Im^2^)/Re, and to estimate the fusion-pore diameter (2*r*) using the equation *G*_p_□=□(π*r*^2^)/(ρλ), where ρ is saline resistivity (100 Ω cm) and λ is the estimated fusion-pore length (15 nm)^100^.

Exocytotic events in Im were selected manually by the cursor option in CellAn (Celica Biomedical, Slovenia), written for MATLAB. An exocytotic event was considered detectable if the signal-to-noise ratio was at least 3:1, and the event was not projected onto the current trace. An event was considered transient (reversible) if a step in Im was followed by a subsequent step of the same amplitude and opposite direction within 15 s, and full (irreversible) in the absence of a reciprocal step. We excluded events with *C*_v_ >10 fF or *G*_p_ >500 pS.

In a subset of electrophysiological experiments, we used transfected astrocytes with knockdown amisyn or overexpressing EGFP-tagged amisyn (wt or PH mutant). These cells were identified via monochromator (Polychrome IV, TILL Photonics) illumination at an excitation wavelength of 488 ± 5 nm, with the fluorescence emission filtered at 515-565 nm and captured with a TILL IMAGO CCD camera. The noise level in *C*_m_ recordings was, on average, slightly higher in transfected cells (82 ± 5 aF RMS, *n* = 171) than in non-transfected cells (59 ± 5 aF RMS, *n* = 211). However, when measured, the *C*_m_ steps were at least 3 standard deviations above the noise.

### Statistical analysis

Unless stated otherwise, all data are displayed as means ± SEM (standard error of the mean). Based on the statistical properties of the analysed data and the number of groups compared (two or more), we used parametric (Student’s t test or one-way ANOVA, respectively) or non-parametric (Mann-Whitney U test or ANOVA on ranks followed by Dunn’s test, respectively) tests to determine the statistical significance. The statistical tests used are specified for each figure. In the analysis of the electrophysiological parameters that were calculated or normalized per recording (event frequency, percentage of projected events), we weighted the mean and variance (by effective time of recording, number of events per recording) to reduce the bias of short or quiet recordings and followed with a respective parametric test. Statistical analysis was performed using SigmaPlot 11.0 (Systat Software, San Jose, CA, USA) and R (R Development Core Team, 2013). The graphics in the figures were prepared using SigmaPlot 11.0 and the ggplot2 library^101^ in R.

## Author contributions

J.V., M.B., Z.B. performed electrophysiological membrane capacitance measurements and analyzed the recordings. J.V. conducted immunocytochemical stainings. J.V., Z.B., M.S. performed optophysiological experiments. J.V., M.S. performed image analysis and quantification, analyzed the data, performed data visualization and constructed figures. K.S, K.D., S.P. carried out molecular biology experiments, including siRNA validation and gene-expression analyses. M.P., Z.B., G.A., I.M. carried out plasmids construction and validation. J.J. assisted with electrophysiological membrane capacitance measurements, data analysis, statistical evaluation, and manuscript writing. M.K. contributed to experimental design and interpretation of the membrane capacitance measurements. M.S. conceived the experiments, designed and performed the confocal imaging experiments and wrote the manuscript. R.Z. generated the idea, conceived the study, directed and supervised the project, secured funding, contributed to experimental design, and wrote and revised the manuscript. All authors discussed the results, critically revised the manuscript and approved the final version.

## Extended Data Material

**Extended Data Fig. 1.**
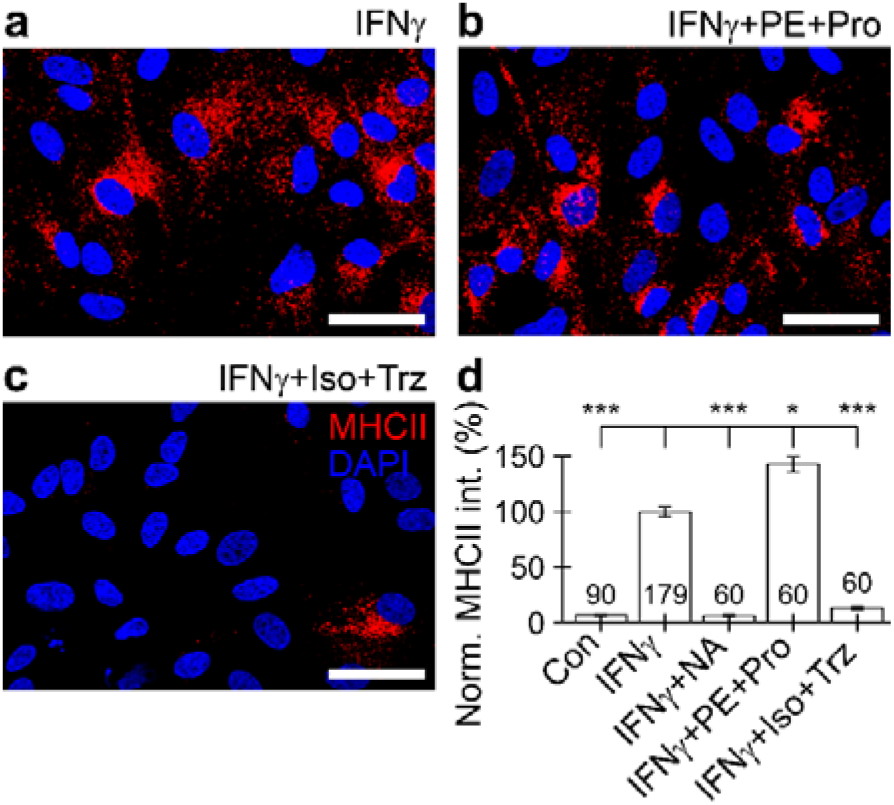
Prolonged treatment with β-adrenergic receptor agonists reduces cellular MHCII expression in IFNγ-treated astrocytes: fixed and permeabilized. (**a–c**) Confocal images of fixed and permeabilized astrocytes immunolabelled by the anti-MHCII antibody and the corresponding fluorescent secondary antibody (MHCII, red), and by the fluorescent nuclear stain DAPI (blue), after 48 h of treatment with 100 U/ml of IFNγ (**a**), co-treatment with 100 U/ml of IFNγ, 100 µM PE, and 2 µM Pro (**b**) and co-treatment with 100 U/ml IFNγ, 100 µM Iso, and 2 µM Trz (**c**). Note the reduction in MHCII immunofluorescence in IFNγ-treated astrocytes co-treated with Iso and Trz. (**d**) Graph displaying mean (±SEM) normalized MHCII fluorescence per astrocyte subjected to different (co)treatments with IFNγ and α-and β-adrenergic (ant)agonists. Numbers at the bottom of the bars indicate the number of images analysed. \**P* < 0.05, \*\*\**P* < 0.001, n.s. not significant (ANOVA on ranks followed by Dunn’s test). Co, control; IFNγ, interferon γ; Iso, isoprenaline; NA, noradrenaline; PE, phenylephrine; Pro, propranolol; Trz, terazosin.

**Extended Data Fig. 2.**
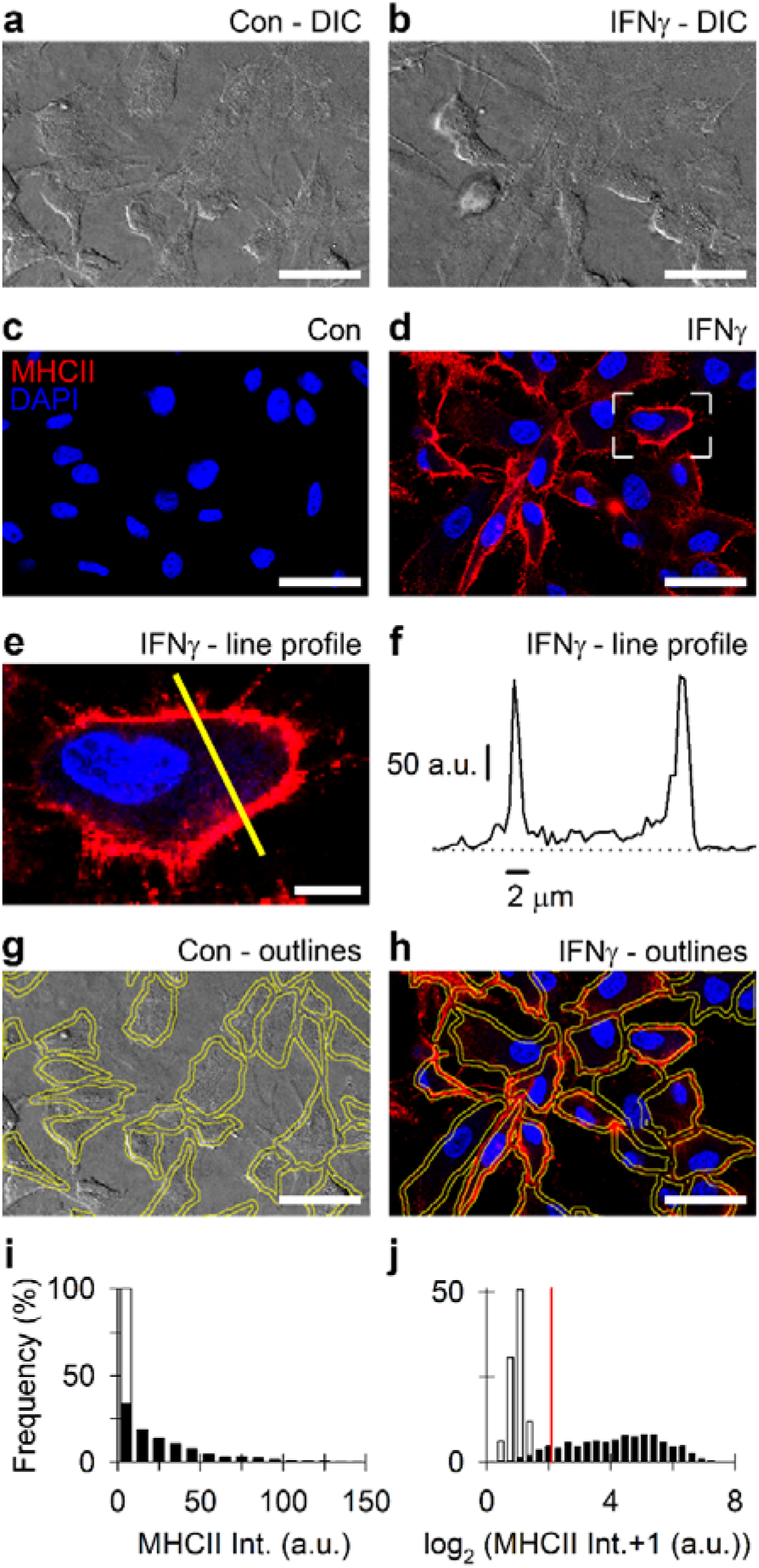
IFNγ treatment induces surface expression of MHCII in cultured rat astrocytes. (**a–d**) Differential interference contrast (**a, b**) and confocal micrographs (**c, d**) of non-permeabilized cultured rat astrocytes not treated (Con; **a, c**) or treated with 60 U/ml of IFNγ for 48 h (**b, d**), immunolabelled with anti-MHCII antibodies and the corresponding fluorescent secondary antibodies (MHCII, red), and with the fluorescent nuclear stain DAPI (blue). Scale bars (**a–d**): 50 µm. (**e**) Magnified view of an IFNγ-treated MHCII-positive (MHCII+) astrocyte selected by the open white frame in (**d**). The thin yellow line indicates the fluorescence intensity profile drawn over the MHCII+ astrocyte. Scale bar: 10 μm. (**f**) The fluorescence intensity line profile (drawn from **e**) indicates MHCII surface expression, as highlighted by distinctive peaks demarcating the cell perimeter. (**g, h**) DIC and confocal micrographs of non-treated controls (**g**) and IFNγ-treated astrocytes (**h**) with regions (outlined in yellow; band thickness 2 µm) used to measure MHCII immunofluorescence at the cell periphery. Scale bars (**g, h**): 50 µm. (**i**) Superimposed frequency histograms displaying MHCII fluorescence intensity at the periphery of non-treated controls (white) and IFNγ-treated astrocytes (black). (**j**) Superimposed frequency plots of logarithmically (log_2_) transformed data from (**i**). The thin vertical red line indicates peripheral MHCII fluorescence of 4 a.u., which is used as the threshold for MHCII+ astrocytes. Con, control; DIC, differential interference contrast; IFNγ, interferon γ.

**Extended Data Fig. 3.**
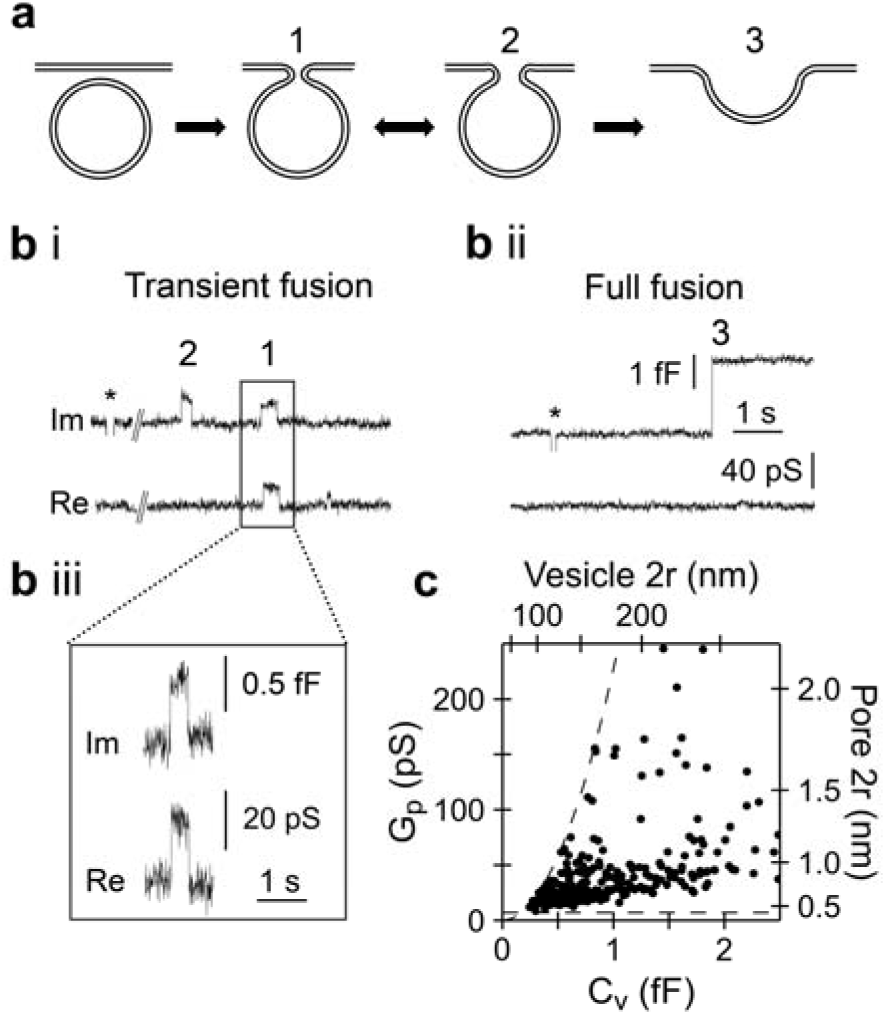
High-resolution cell-attached membrane capacitance measurements of single-vesicle exocytosis. (**a**) Stages of vesicle exocytosis. The vesicle docks and fuses with the plasmalemma to establish a narrow fusion pore (1), which then widens reversibly (2) or irreversibly to favour merger of the vesicle membrane with the plasmalemma (3). (**b**) Discrete upward and downward steps in the imaginary (Im) and real (Re) components of the admittance signal correspond to elementary events of: (**i**) transient fusion, (**ii**) full fusion, (**iii**) transient fusion (box in **i**) exhibiting projection to the Re trace, indicating the formation of a highly resistive fusion pore enabling measurement of fusion-pore conductance (*G*_p_). The 0.53 fF (Im) and 19.8 pS (Re) steps correspond to a vesicle with capacitance (*C*_v_) of 0.99 fF and *G*_p_ of 42.5 pS, translating to a vesicle diameter of 180 nm and pore diameter of 0.9 nm, respectively. Asterisks denote calibration pulses. (**c**) Scatter plot depicts the relationship between *C*_v_ and *G*_p_ in exocytotic vesicles (*n* = 655). Larger vesicles tend to establish fusion pores with larger diameters. Dashed lines indicate detection limits for *G*_p_ projections*, at a minimal signal-to-noise ratio of 3 and assuming the intrinsic noise of 0.06 fF (2.4 pS) RMS. The exocytosing vesicles without projections (not shown) were smaller (0.40 ± 0.02 fF; 107 ± 2 nm; *n* = 302) than vesicles with projections (1.13 ± 0.05 fF; 177 ± 4 nm; *n* = 353). *Debus, K. & Lindau, M. Resolution of patch capacitance recordings and of fusion-pore conductances in small vesicles. *Biophys J* **78**, 2983-2997 (2000).

**Extended Data Fig. 4.**
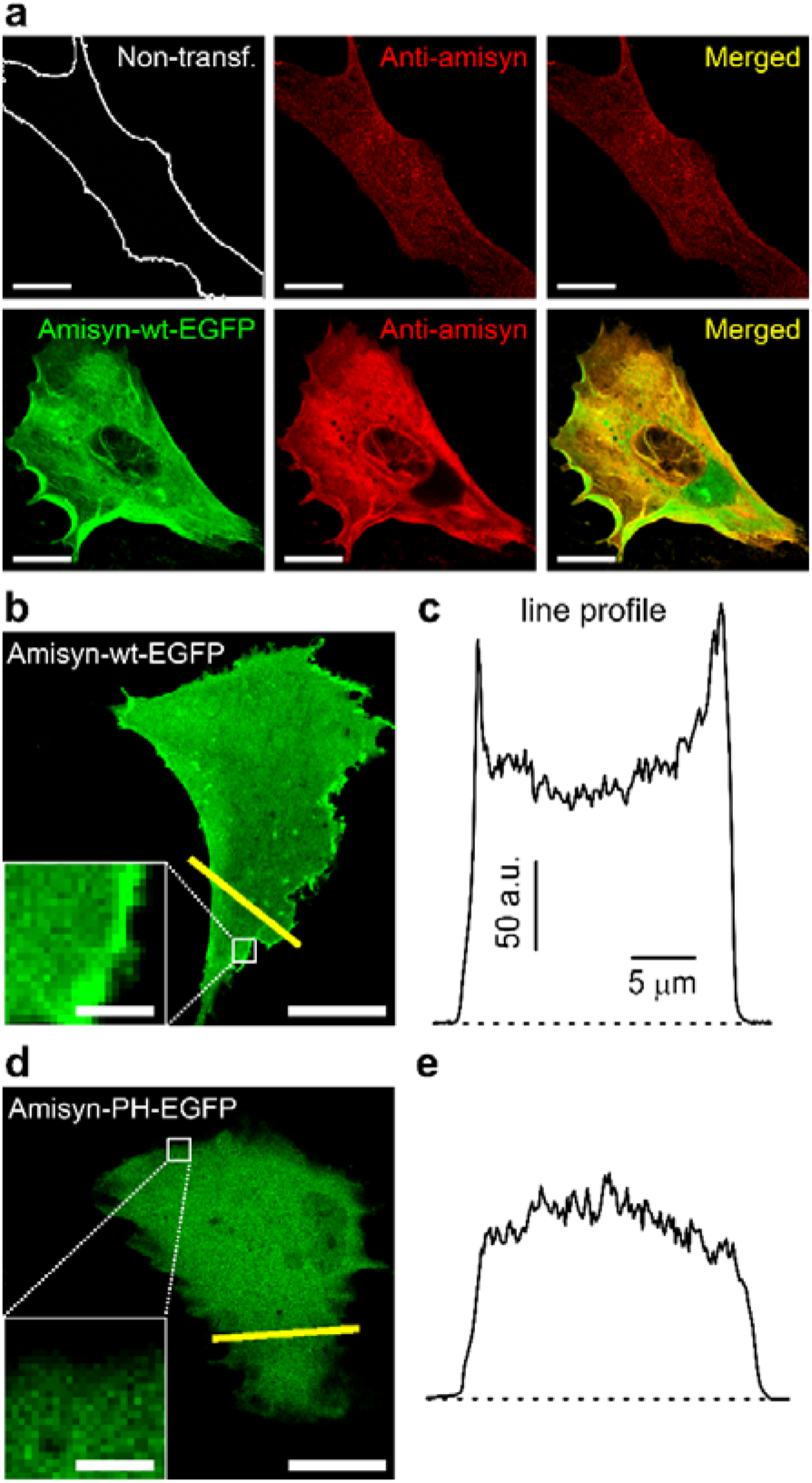
Amisyn is natively expressed by cultured rat astrocytes. (**a**) Confocal micrographs of non-transfected (top left) and transfected astrocytes (bottom left) overexpressing amisyn-wt-EGFP (green, left) that were immunolabelled with the anti-amisyn antibody and with the corresponding fluorescent secondary antibody (red, middle). The merged images display co-localized fluorescence (yellow, right). Note the stronger anti-amisyn immunofluorescence in transfected astrocytes. Scale bars: 20 μm. EGFP, enhanced green fluorescent protein; wt, wild-type.Amisyn’s association with the astrocyte plasmalemma depends on an intact pleckstrin homology domain. (**b, e**) Confocal micrographs of cultured astrocytes overexpressing amisyn-wt-EGFP (**b**) and amisyn-PH-EGFP (**d**). Note the enriched plasmalemmal localization of amisyn-wt, but not amisyn-PH, visually indicated by the bright fluorescent edge contrasting the dimmer cell interior. Insets display a magnified view of the selected regions (white frame) to emphasize the relative enrichment of amisyn-wt-EGFP at the plasmalemma. Scale bars: 20 µm (large images) and 2 μm (insets). (**c, e**) Fluorescence intensity line profiles drawn across astrocytes in (**b**) and (**d**) (yellow lines). Note fluorescence peaks at the edge of the line profile in astrocytes expressing amisyn-wt-EGFP, but not amisyn-PH-EGFP. Fluorescence peaks suggest amisyn-wt association with the plasmalemma, whereas the absence of fluorescence peaks indicates that mutated amisyn (point mutations in the PH domain: K30A, K32A, K64D, K66D) was unable to associate with the plasmalemma and therefore distributed only in the cell cytosol. EGFP, enhanced green fluorescent protein; PH, pleckstrin homology; wt, wild-type.

**Extended Data Fig. 5.**
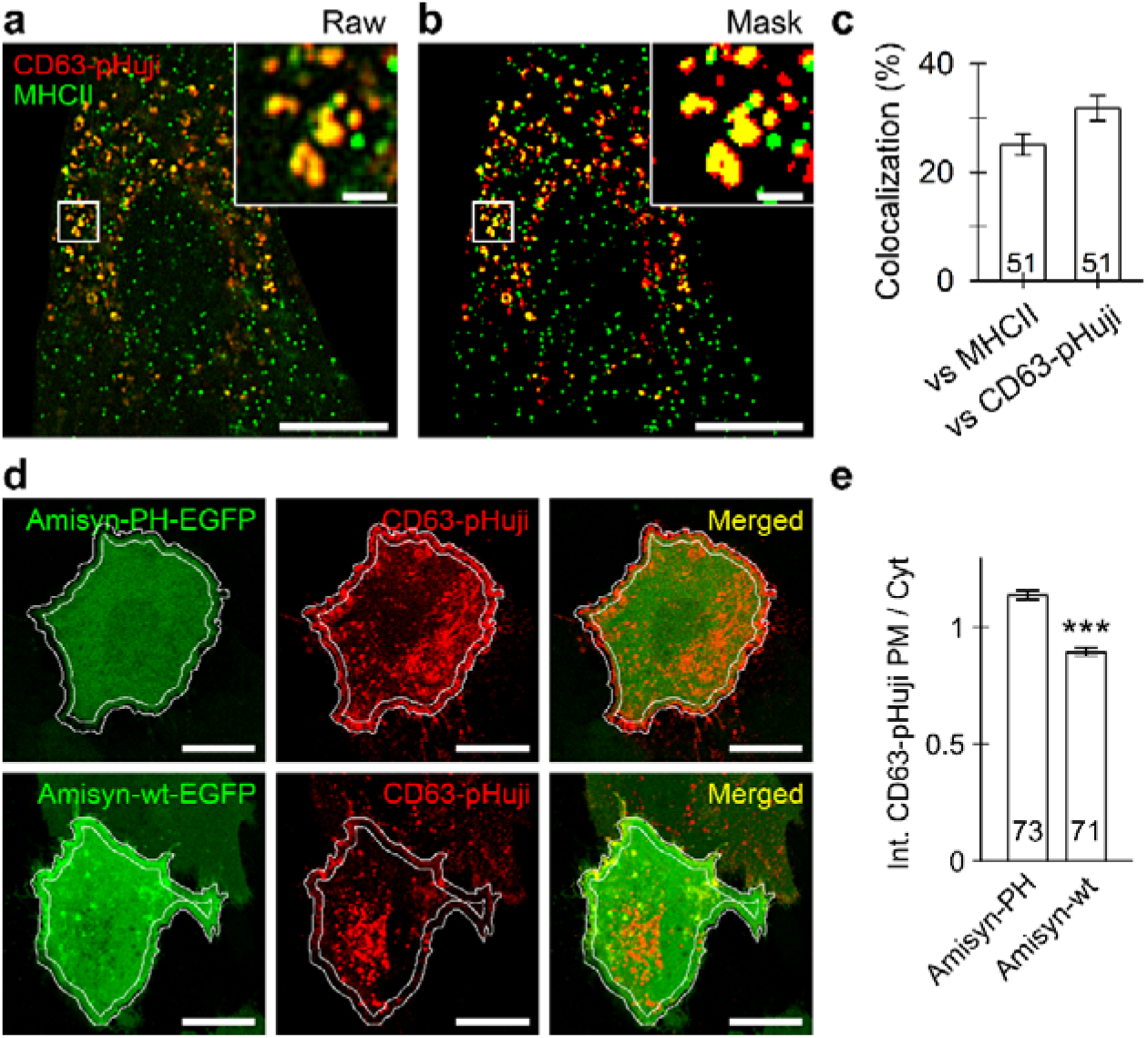
Overexpression of wild-type amisyn reduces surface expression of CD63-pHuji in astrocytes. MHCII localizes to the same vesicles as the lysosomal membrane protein CD63. (**a**) Super-resolution structured illumination microscopy (SIM) micrographs of fixed, double-fluorescent IFNγ-treated astrocytes display immunofluorescent MHCII (green) and CD63-pHuji-positive compartments (red). (**b**) The thresholded mask image (from **a**) shows co-localized pixels (yellow); the insets show a magnified view of the selected vesicles (within the white frame). Scale bars: 10 μm (large images) and 1 μm (insets) (**a, b**). (**c**) Graph displaying quantitative co-localization (mean ± SEM) of CD63-pHuji versus anti-MHCII fluorescence and vice versa. Numbers at the bottom of the bars indicate the number of cell images analysed. IFNγ, interferon γ. (**d**) Confocal micrographs of live double-transfected astrocytes expressing amisyn-PH-EGFP or amisyn-wt-EGFP (green, left) and CD63-pHuji-positive compartments (red, middle). The merged images display co-localized fluorescence (yellow, right). The white outline marks a 2-µm band at the plasmalemma used to measure CD63-pHuji fluorescence; the area below the band represents cytosolic fluorescence. Scale bars: 20 μm. (**e**) Graph displaying the intensity ratio (mean ± SEM) of CD63-pHuji at the plasmalemma and in the cytoplasm in astrocytes expressing amisyn-PH-EGFP or amisyn-wt-EGFP, respectively (see the Image analysis section in the Methods). Numbers at the bottom of the bars indicate the number of cell images analysed. \*\*\**P* < 0.001 (Mann-Whitney U test). Cyt, cytoplasm; EGFP, enhanced green fluorescent protein; PH, pleckstrin homology; PM, plasmalemma, wt, wild-type.

**Extended Data Fig. 6.**
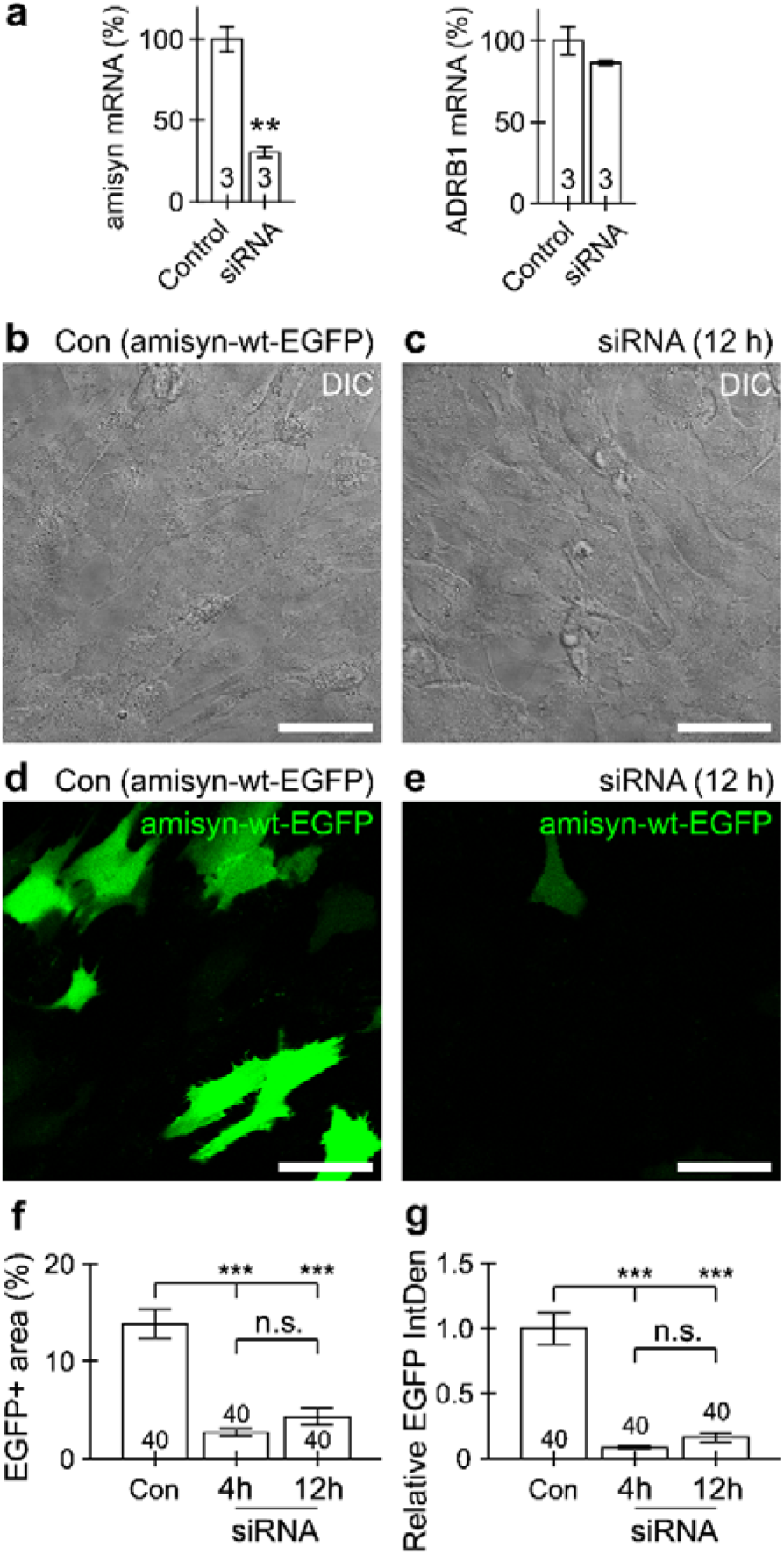
Astrocyte co-transfection with amisyn siRNA (knockdown) and pAmisyn-wt-EGFP decreases the expression of amisyn-wt-EGFP. (**a**) siRNA-mediated knockdown of amisyn specifically reduces amisyn mRNA levels without affecting adrenoreceptor β1 (ADRB1) expression. Cortical astrocytes were mock-transfected (Control) or transfected with siRNA targeting amisyn (siRNA) and then analysed for expression of amisyn (left) and ADRB1 (right) mRNA with quantitative polymerase chain reaction. Data represent mean ± SEM (*n* = 3). \*\**P* < 0.01 versus control; Student’s *t* test. mRNA, messenger RNA; siRNA, small interfering RNA. (**b, c**) Differential contrast micrographs of astrocytes indicate similar cell density in the two cultures; (**d, e**) confocal micrographs of the same astrocytes expressing amisyn-wt-EGFP indicate a profound reduction in the number of amisyn-wt-EGFP-expressing astrocytes when cells were co-transfected with siRNA (**e**) compared with amisyn-wt-EGFP-expressing controls (**d**). (**f, g**) Quantitative analysis of amisyn-EGFP-expression after different siRNA exposure conditions (change of culture media after 4 h or 12 h, in both cases imaged after ∼24 h). Graphs depict the relative area (in %; mean ± SEM) of the amisyn-wt-EGFP signal in the image (**f**), which indicates the number of amisyn-wt-EGFP-positive cells, and the relative integrated density of amisyn-wt-EGFP fluorescence within the image (**g**). Numbers at the bottom of the bars depict the number of images analysed. \*\*\**P* < 0.001, n.s. not significant (ANOVA on ranks followed by Dunn’s test). Based on these results, we can estimate >80% reduction in the number of transfected cells and >90% reduction in total amisyn-wt-EGFP expression. Con, control; DIC, differential interference contrast; EGFP, enhanced green fluorescent protein; IntDen, integrated density; siRNA, small interfering RNA; wt, wild-type.

**Extended Data Fig. 7.**
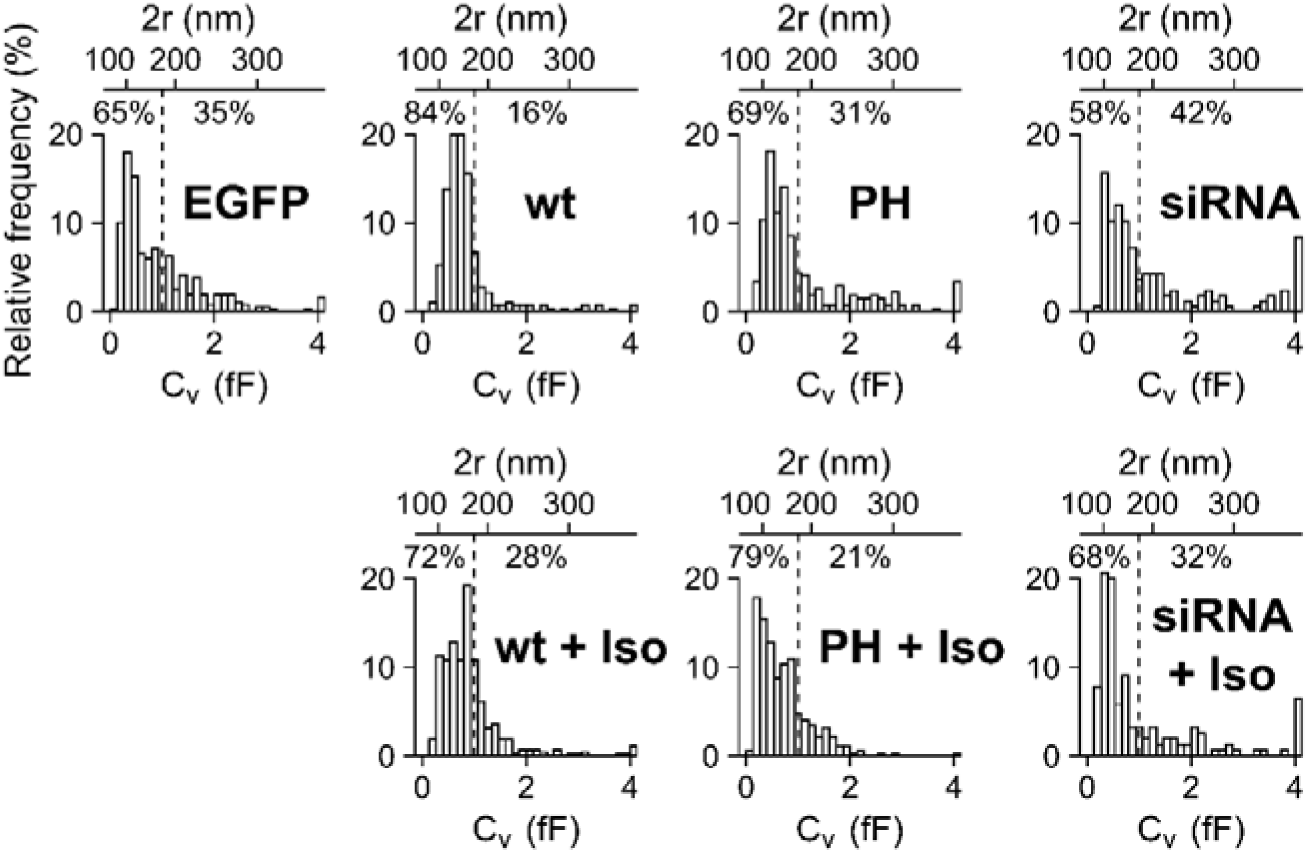
Differential expression of amisyn affects transient exocytosis of larger vesicles. Relative frequency distributions of vesicle capacitance (*C*_v_, bottom) and vesicle diameter (top) of transient exocytotic events in all groups examined (described in the legend to Fig. 6). The dashed vertical line delimits the percentage of vesicles smaller or larger than 1 fF (corresponding to a vesicle diameter of ∼178 nm). Note the increase in the proportion of largest transient exocytotic vesicles (>4 fF; corresponding to a vesicle diameter of >357 nm) in astrocytes with knockdown of amisyn (siRNA, siRNA + Iso). Also note the decreased proportion of larger transient exocytotic vesicles (>1 fF) in astrocytes with overexpressed amisyn and/or after β-adrenergic treatment (wt, wt + Iso, PH + Iso). EGFP, enhanced green fluorescent protein; Iso, isoprenaline; PH, pleckstrin homology; siRNA, small interfering RNA; wt, wild-type.

**Extended Data Fig. 8.**
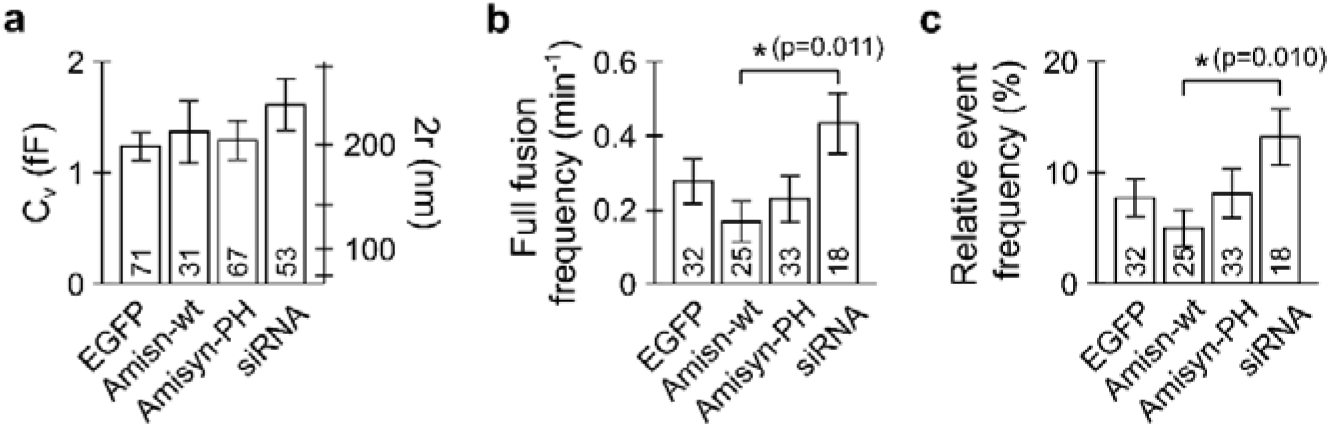
Amisyn decreases the probability of full fusion. The effects of amisyn variant overexpression or knockdown on full-fusion events. (**a–c**) Graphs showing (mean ± SEM) vesicle capacitance (*C*_v_; **a**), frequency of full-fusion events (**b**), frequency of full-fusion events relative to all fusion events (**c**). Numbers at the bottom of the bars denote the number of events (**a**) or the number of recordings (**b, c**) analysed. t test \**P* < 0.05. EGFP, enhanced green fluorescent protein; PH, pleckstrin homology; siRNA, small interfering RNA; wt, wild-type

